# Pseudocobalamin production and use in marine *Synechococcus* cultures and communities

**DOI:** 10.1101/2024.04.01.587587

**Authors:** Catherine C. Bannon, Maria A. Soto, Elden Rowland, Nan Chen, Anna Gleason, Emmanuel Devred, Julie LaRoche, Erin M. Bertrand

## Abstract

Cobalamin influences marine microbial communities because an exogenous source is required by most eukaryotic phytoplankton, and demand can exceed supply. Pseudocobalamin is a cobalamin analog that is produced and used by most cyanobacteria but is not directly available to eukaryotic phytoplankton. Some microbes can remodel pseudocobalamin into cobalamin, but a scarcity of pseudocobalamin measurements impedes our ability to evaluate its importance for marine cobalamin production. Here, we perform simultaneous measurements of pseudocobalamin and methionine synthase (MetH), the key protein that uses it as a co-factor, in *Synechococcus* cultures and communities. In *Synechococcus* sp. WH8102, pseudocobalamin quota decreases in low temperature (17 °C) and low N:P, while MetH did not. Pseudocobalamin and MetH quotas were influenced by culture methods and growth phase. Despite the variability present in cultures, we found a comparably consistent quota of 300 ± 100 pseudocobalamin molecules per cyanobacterial cell in the Northwest Atlantic Ocean, suggesting that cyanobacterial cell counts may be sufficient to estimate pseudocobalamin inventories in this region. This work offers insights into cellular pseudocobalamin metabolism, and the environmental and physiological conditions that may influence it, and provides environmental measurements to further our understanding of when and how pseudocobalamin can influence marine microbial communities.

## Introduction

Cobalamin, B_12_, is an essential organic micronutrient that impacts microbial community composition and productivity in various parts of the ocean (Koch *et al*., 2011; Bertrand *et al*., 2015). Most cyanobacteria produce and use a unique cobalamin analog called pseudocobalamin, psB_12_, that is poorly bioavailable to most eukaryotic phytoplankton because of its structural differences at the α-ligand (adenine = psB_12_, DMB = B_12_) (Helliwell *et al*., 2016; Heal *et al*., 2017). Similar to B_12_, psB_12_ occurs in different forms (Me-, Ado- and OH-psB_12_), which have unique functions in the cell (Fig 1A). psB_12_ has the potential to impact B_12_ availability through the activities of remodelers, such as select heterotrophic bacteria and eukaryotic algae, which are able to exchange the adenine α-ligand for DMB (Helliwell *et al*., 2016; Ma *et al*., 2020). The prevalence of and controls on psB_12_ production and use remains unclear, in part due to the limited measurements of psB_12_ in cultures and in the natural environment.

**Figure 1:**
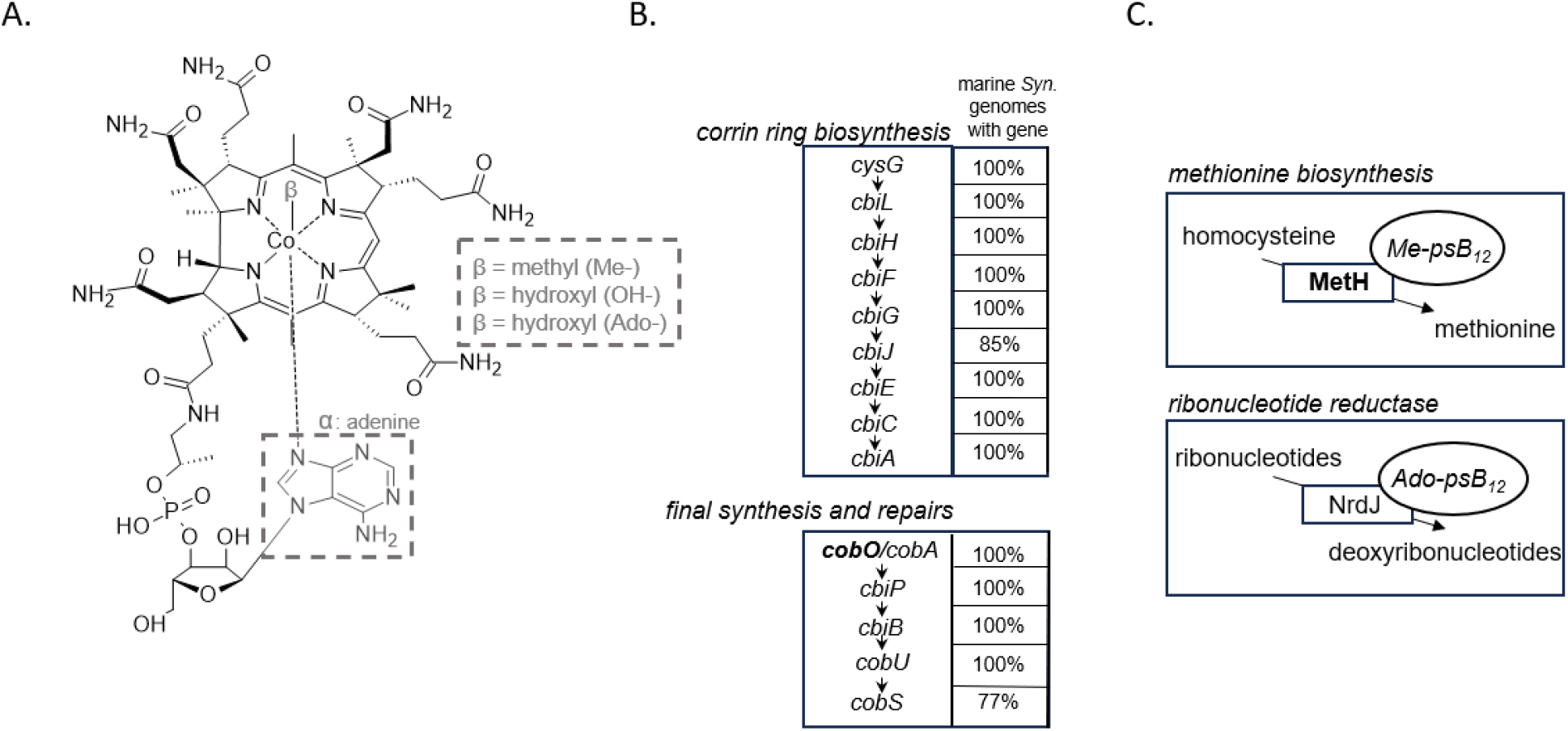
(A) Structure of pseudocobalamin, psB_12_, with adenine at the alpha (α) ligand and Me, OH- or Ado- at the beta (β) ligand. C:N ratios of the molecules are as follows (OH-psB_12_ = 3.6:1, Me-psB_12_ = 3.6:1, Ado-psB_12_ = 3.2:1). (B) Simplified diagram of cobalamin biosynthesis pathway along with the percent of marine *Synechococcus* genomes (n = 15) with the putative gene from Helliwell et al, 2016. (C) Known (MetH) and expected (NrdJ) proteins in *Synechococcus* that use psB_12_ as a co-factor. Genes/proteins that are bolded are investigated in this study as proxy.

Accurately quantifying metabolites can help characterize the composition, activity, and controlling processes in marine microbial communities. However, such measurements remain difficult in part because of the complex and unknown chemical backgrounds they are embedded in (Kido Soule *et al*., 2015; Moran *et al*., 2022). Multiple studies have quantified particulate B_12_ (size fraction >0.2 µm), while only one study to date has quantified particulate psB_12_ in the ocean (Heal *et al*., 2017). Concentrations of particulate psB_12_ ranged from undetectable to 0.15 pM. These values correlated with cyanobacterial carbon and co-occurred with B_12_ (Heal *et al*., 2017). Quantifying both psB_12_ and B_12_ compounds would provide a more complete picture of the processes governing cobalamin availability, especially in regions vulnerable to cobalamin co-limitation and where cyanobacteria are abundant.

Insight into psB_12_ cell biology is restricted because psB_12_ quotas have only been quantified to date in four cyanobacterial strains (Heal *et al*., 2017) and the influence specific biotic or abiotic factors have on psB_12_ production and use have yet to be explored, either in culture or in the field. The psB_12_ biosynthesis pathway includes up to 30 enzymes (Cob/Cbi proteins) and is similar to the B_12_ pathway with the exception of the lower ligand synthesis and activation (DMB for B_12_, adenine for psB_12_) (Helliwell et al, 2016; Roth et al, 2003). Corrin ring biosynthesis is present in the majority of available marine *Synechococcus* genomes (n = 15) (Helliwell et al, 2016) (Fig 1B) but the specific bottleneck or rate limiting step of this pathway remains unclear (Rodionov *et al*., 2003; Lu *et al*., 2020). One key psB_12_-dependent enzyme in cyanobacteria is methionine synthase (MetH or MS), which requires *methyl*-psB_12_ (Me-psB_12_) as a cofactor to generate methionine from homocysteine and 5-methyltrahydrofolate (Tanioka *et al*., 2009, 2010; Heal *et al*., 2017) (Fig 1C). An alternative non-B_12_ requiring methionine synthase, MetE, appears to be absent in marine *Synechococcus* genomes (Rodinov et al, 2003). Some cyanobacteria also encode NrdJ, a class II ribonucleotide reductase that is expected to use Ado-psB_12_ to generate dNTPs from their ribonucleotide metabolites through a deoxygenation step (Gleason and Olszewski, 2002; Lundin *et al*., 2010), though the use of psB_12_ by this enzyme has yet to be specifically examined (Fig 1C). Quantifying proteins involved in both psB_12_ production and use has the potential to provide insights into psB_12_ cell biology, but has thus far not been investigated.

*Synechococcus* is one of the most abundant cyanobacterial genera in the ocean (Flombaum *et al*., 2013) and an important member of the microbial community in the Northwest Atlantic Ocean, particularly in coastal euphotic zone waters during the fall (Li and Wood, 1988; Zorz et al, 2019; Soto et al, 2023), As such, it is likely to be a major source of psB_12_ in the region. *Synechococcus* is comprised of many strains that can be grouped into distinct clades using phylogenetic clustering (Ahlgren and Rocap, 2012). These clades appear to thrive under distinct levels of light, temperature, macronutrients, and metals (Sohm et al, 2016; Ahlgren and Rocap, 2012). Environmental trends in this region indicate that the plankton community is subject to large changes in temperature (from 2 °C in spring up to 25 °C in fall) (Zorz et al, 2019) and nitrogen availability (Loder *et al*., 2013). Dominant *Synechococcus* ASVs in the Northwest Atlantic Ocean have been placed in clade III, I and IV (Robicheau *et al*., 2022). Investigating the influence that temperature and nutrient availability have on psB_12_ in a *Synechococcus* isolate of a regionally important clade could provide insight into how the psB_12_ inventory may fluctuate in this region.

Here, we leverage targeted metabolomics and proteomics to investigate psB_12_ production and use in *Synechococcus* in communities from the Northwest Atlantic Ocean (NWA). To begin, we cultured *Synechococcus* sp. WH8102, a well characterized strain belonging to a dominant clade in the region (clade III), under a range of conditions (low N:P, low temperature, diel cycle, growth phase) and distinct culture methods (batch culture, semicontinuous culture). By measuring both psB_12_ production and use in these conditions, we obtain quantitative insights into *Synechococcus* psB_12_ cell biology while constraining psB_12_ quotas and highlighting environmental and physiological conditions that may impact it. Then, using samples collected from the Northwest Atlantic Ocean, we investigate in situ psB_12_ quotas and use in environmental *Synechococcus* communities through protein and co-factor quantification.

## Material and methods

### Identification of proteins and selection of peptides

We performed a literature search to identify proteins involved in psB_12_ production (Cob/Cbi proteins) and utilization (MetH and NrdJ). BLASTp was used to search for homologues of these *Synechococcus* sp. WH8102 proteins in other available marine *Synechococcus* genomes using the GenBank nr database. Proteins were aligned using COBALT (Papadopoulos and Agarwala, 2007) and digested in silico with trypsin to reveal conserved tryptic peptides. Sequences were introduced in the Unipept Tryptic Peptide Analysis search (Singh *et al*., 2019) to determine if the peptide is present in a significant number of the *Synechococcus* strains of interest. Peptides for CobO, cob(I)alamin adenosyl transferase, and MetH, methionine synthase, were selected for analyses as they met the peptide selection guidelines for SRM targeted assays described in Hoofnagle et al. (2016) (Hoofnagle *et al*., 2016) and were largely conserved across *Synechococcus* clades. Peptides are reported in Table S1. We found no suitable peptide with adequate coverage across *Synechococcus* clades for NrdJ.

### Culture experiments to measure *Synechococcus* peptides and pseudocobalamin quotas

*Synechococcus* sp. WH8102 (CCMP2370) obtained from the National Center for Marine Algae and Microbiota (NCMA, Bigelow Laboratory, USA) was grown axenically in plastic, vented 250 mL flasks (10861-576, VWR) with modified Synthetic Ocean Water media (Dupont *et al*., 2008) void of any added cobalamin, under ∼20 μmol photons m^−2^ s^−1^ supplied through white LED lights (LEDMO-EZ550). Irradiance was measured by a PAR irradiance sensor (QSL2101, Biospherical Instruments) and set at a light:dark cycle of 12:12 h (Table 1). Throughout culturing, we monitored growth via relative fluorescence units (RFU) measured in a 10-AU Fluorometer (Turner Designs, San Jose, CA), and cell counts using a BD Accuri™ C6 flow cytometer as described below. Cultures were also tested for axenicity once a week using SYBR Green II dye (Invitrogen) followed by flow cytometry screening. Specific growth rates (d^−1^) were calculated using linear regression of ln cell counts during exponential growth.

**Table 1:** Experimental treatments (n=3) performed on *Synechococcus* sp. WH8102 in specific culture methods.

| Treatment | Culture methods | Harvesting time |
| --- | --- | --- |
| Control ($24 \pm 1$ °C; 26 N:P) | Semicontinuous | after ~7 generations<br>(Exponential phase) |
| Low temperature ($17 \pm 1$ °C) | | |
| Low N:P ratio (10 N:P) |  |  |
| Diel cycle (harvest at 9:30,<br>18:00 and 22:00) | Batch | Exponential phase |
|  |  | Stationary phase |

We maintained semicontinuous cultures of *Synechococcus* sp. WH8102 in exponential phase by replacing a volume of culture with fresh media to dilute the cell concentration to a fixed value every other day (Table 1). Semicontinuous cultures were grown in three conditions, control (24±1 ºC; 26 N:P), low temperature (17±1 ºC; 26 N:P) and low N:P ratio (24±1 ºC;10 N:P). Cultures (n=3) were maintained for ∼7 generations in exponential phase under the conditions described before gentle harvesting using a bottle top vacuum filtration system. We harvested protein samples (50 mL) on 0.2 µm polycarbonate filters, metabolite samples (50 mL) on 0.2 µm Nylon filters, and particulate organic carbon (POC) samples (20 mL) on GF/F glass microfiber filters.

For strain *Synechococcus* sp. WH8102 grown in batch culture, we harvested samples at both exponential and stationary phase (Fig S1). For early exponential phase, we harvested 35 mL for protein samples, 17 mL for metabolite samples, and 8 mL for POC samples at the three points in the diel cycle. In stationary phase, we harvested 20 mL for protein samples, 10 mL for metabolite samples, and 5 mL for POC samples at the three points in the diel cycle. The three points of the diel cycle evaluated were: 1.5 hours after the lights went on (9:30), 1 hour before the lights went off (18:00), and 3 hours after the lights went off (22:00).

### *Synechococcus* sp. WH8102 culture cell counts

Culture cell density was measured on an Accuri C6 flow cytometer (BD Biosciences) equipped with a Red (640 nm) and Blue (488 nm) laser, after labeling with1:1000 SYBR Green II dye (Invitrogen). Single cell events were recorded on forward side scatter and side scatter plots and distinguished using optical filters of phycoerythrin (488 nm laser, filter = 572/28 nm) and chlorophyll *a* (488 nm laser, filter = 675/30 nm). Accuri run limits were set to a maximum of 500,000 events or 50 µL, with a flow rate of 35 µL/min and a core size of 16 µm. The gating strategy for *Synechococcus* populations was developed based on the Acccuri application note provided by BD Biosciences.

### POC measurements

Samples were acidified for 6 hours in a glass desiccator with an open bottle of concentrated hydrochloric acid (37% HCl, 250 mL) and later dried overnight at 45 °C. Filters were packed in tin capsules and analyzed for particulate carbon on an elemental analyzer (Elementar Vario microcube) coupled to an IRMS (Isoprime 100). The samples were flash combusted at 1150 °C to convert particulate carbon into CO_2_ gas. These gaseous components were then analyzed by the Isoprime 100. The values were blank-corrected and divided by cell number to give per cell quota.

### Environmental sample collection

We collected environmental samples in Fall 2016 onboard an AZMP (Atlantic Zone Monitoring Program) cruise aboard CCGS Hudson from September 15th, 2016, to October 6th, 2016 (HUDSON 2016027). Samples were taken at 4 stations on the Halifax line (HL 01, HL 02, HL 05.5, HL 06). At each of the stations, we collected 1 L of water at each depth for particulate metabolite analysis. The sample was gently filtered through a 0.2 μm Nylon filter in the dark to reduce photodegradation. For protein samples, 10 L of water was pre-filtered (330 μm) from Niskin bottles then filtered gentled through 3 and 0.2 μm polycarbonate filters via peristaltic pumping. Parallel 1.8 mL seawater samples for analytical flow cytometry were fixed with 1% paraformaldehyde. Flow cytometry samples and filters were then frozen immediately at -80°C until analysis.

### Environmental cyanobacterial cell counts

Preserved samples were thawed at room temperature, and prefiltered using 35 μm cell strainers into a 5 mL fluorescence-activated Cell Sorting (FACS) tube. The samples were then collected and analyzed on a Novocyte 3000 with NovoExpress software (Agilent, USA) for natural fluorescence cell counts. The Novocyte 3000 is equipped with three lasers [Red (640 nm), Blue (488 nm), and Violet (405 nm)], and the optical filters of interest are chlorophyll-a (488 nm laser, filter = 675/30 nm),divinyl chlorophyll a (405 nm laser, filter = 675/30 nm), phycocyanin (640 nm, filter =675/30 nm) and phycoerythrin (488 nm laser, filter = 572/28 nm). The samples were collected with a flow rate of 120 μL/min and a threshold of chlorophyll > 450 nm. The gating strategy was developed based on the Novocyte application note provided by Agilent and the *Prochlorococcus* gate was constructed with the aid of pure *Prochlorococcus* culture, an example of environmental flow cytogram is provided (Fig S2).

### Particulate metabolites extraction

Metabolites were extracted from both culture and environmental samples following the methodology described by Heal et al. (2017) with minor modifications. Briefly, 0.2 mL of each 100 µm and 400 µm silica beads were added to the bead beater tubes containing sample filters. 1 mL of ice-cold solvent mixture (40:20:20 acetonitrile: methanol: MQ water) was added and the mixture was agitated using a bead beater (MP Biomedicals) in 3 × 40 second pulses at 1800 rotations per minute (RPM) over a 20-minute period. Heavy-CN-B_12_ (Cambridge Isotopes Labs, CLM-9770-E) was used as an internal standard and was added to a final concentration of 3 nM in all metabolite samples prior to extraction. Solvent was removed under vacuum (Eppendorf, Mississauga, ON) at room temperature. Culture samples were re-suspended in buffer A (20 mM ammonium formate, 0.1% formic acid, 2% acetonitrile) to generate equivalent of extracts from 12,100 cells/µL. Environmental samples were resuspended with 100 µL of buffer A. All samples were then vortexed and centrifuged at 10,000 x g for 3 min at 4 ºC and then diluted 2-fold with buffer in conical polypropylene HPLC vials (Phenomenex, Torrance, CA) prior to mass spectrometry analysis.

### Metabolite quantification

HPLC-MS was used to quantify cobalamins using a Dionex Ultimate-3000 LC system coupled to the electrospray ionization source of a TSQ Quantiva triple-stage quadrupole mass spectrometer in SRM mode. Details of mass spectrometry conditions are reported in supplement methods and the transition list is reported in Table S2.

For metabolite samples, quality control (QC) samples were prepared by mixing equal volumes of each sample within its respective group (1 QC for culture samples and 1 QC for environmental samples). Calibration curves with authentic cobalamin standards were prepared using the QC sample as a matrix and triplicate injections were performed for 0, 0.1, 5, 10, 50, and 100 fmol of each authentic cobalamin standard on analytical column with linearity up to 1000 fmol. All cobalamins were normalized by heavy-CN-B_12_ internal standard to account for variability introduced by matrix effects, during extraction and instrument analysis. Limits of quantitation and limits of detection were calculated as 10x and 3x the variation in the QC sample without spike, respectively and recorded in Table S3.

Given that commercial psB_12_ standards are not available, the following assumptions were made for psB_12_ quantification using associated cobalamin analog standards: (1) B_12_ and psB_12_ ionize with the same efficiency, (2) B_12_ and psB_12_ fragments are generated in a similar fashion, (3) B_12_ and psB_12_’s different elution times do not greatly affect the response.

### Protein extraction and digestion

Protein sample filters were submerged in 750 µL of SDS extraction buffer (2% SDS, 0.1 M Tris/HCl pH 7.5, 5% glycerol, 5 mM EDTA) and incubated for 10 min on ice before being heated for 15 min at 95 ºC in a ThermoMixer C (Eppendorf, Mississauga, ON) at 350 RPM. Samples were then sonicated on ice for 1 min with a Q125 Sonicator (Qsonica Sonicators, Newton, CT) at 50% amplitude and 125 W (pulse 15 s ON, 15 s OFF) then kept at room temperature for 30 min and vortexed every 10 min. The filter was then removed, and samples were centrifuged at 15,000 x g for 30 min at room temperature. 4x volume of ice-cold acetone was added to the supernatant for overnight precipitation of proteins at -20 ºC before washing with 3 × 400 µL of ice-cold acetone and 1 × 400 µL with ice-cold methanol. Following each wash, samples were centrifuged at 15,000 x g for 30 min at room temperature and the supernatant was removed. Pellet was dried down under vacuum (Eppendorf, Mississauga, ON) for 15 min at room temperature. Extracted protein was resuspended in 20 µL of 8 M urea for 10 min at room temperature then80 µL of freshly made 50 mM ammonium bicarbonate was gradually added to get a final solution of 1.6 M urea and 40 mM ammonium bicarbonate.

Aliquots of 50 µg of protein were removed from each sample and diluted to 100 µL with 50 mM ammonium bicarbonate for protein digestion. Proteins were reduced with 5mM dithiothreitol (DTT) at 56 ºC for 30 min in a ThermoMixer, then alkylated with 15 mM iodoacetamide at room temperature in the dark. Residual iodoacetamide was quenched with a second addition of DTT to give a final concentration of 10 mM DTT. Protein was digested with 1 µg trypsin (Thermo Scientific, Waltham, MA) at 37 ºC overnight. 0.5 µL of formic acid was added to halt the digestion. A modified protein Micro BCA kit (Thermo Scientific - p/n 23235)was conducted to confirm protein concentration of each sample before injection using a trypsin digested BSA (Bovine Serum Albumin) standard to make the calibration curve.

### Peptide quantification

Targeted mass spectrometry was performed using a Dionex Ultimate 3000 UPLC system interfaced to a TSQ Quantiva triple-stage quadrupole mass spectrometer (MS) (Thermo Scientific, Waltham, MA), fitted with a heated, low flow capillary ESI probe (HESI-II). Isotopically labeled, heavy internal standard versions of each peptide were synthesized by Thermo Scientific™ at >95% purity. Heavy peptide standard was added to samples to yield 20 fmol on column for every 1 ug total protein injected. Details of mass spectrometer conditions are reported in supplement methods. Each sample was analyzed via triplicate injections using the transition list of peptides that is found in Table S1.

### Statistical analysis

Statistical analyses were performed using R (RStudio Team, 2020). A one-way ANOVA followed by a Post-hoc Tuckey’s Test was used to determine the differences in *Synechococcus* culture experiments. A two-way ANOVA followed by a Post-hoc Tuckey’s Test was used to determine the effects of growth phase and diel cycle on the same variables. Differences were considered significant when a p-value < 0.05 was observed.

## Results

### Factors that influence psB_12_ production and use in *Synechococcus* sp. WH8102

We performed culture experiments with *Synechococcus* sp. WH8102 (clade III) in triplicate (n=3) to investigate the influence different conditions have on psB_12_ production and use. Growth rates (per day) were similar to those reported in literature and were consistent across batch grown cultures and semi-continuously grown cultures in control and low nitrogen treatments but significantly lower in low temperature (17°C) (Fig 2A) (Mackey *et al*., 2013). Mol carbon per *Synechococcus* cell increased during the day and in stationary phase, and significantly increased in low temperatures (Fig 2 B, C). Microgram (µg) protein per *Synechococcus* cell was invariant across the time of day and growth phase, but significantly increased in low temperature compared to other treatments (Fig S3B, C).

**Figure 2:**
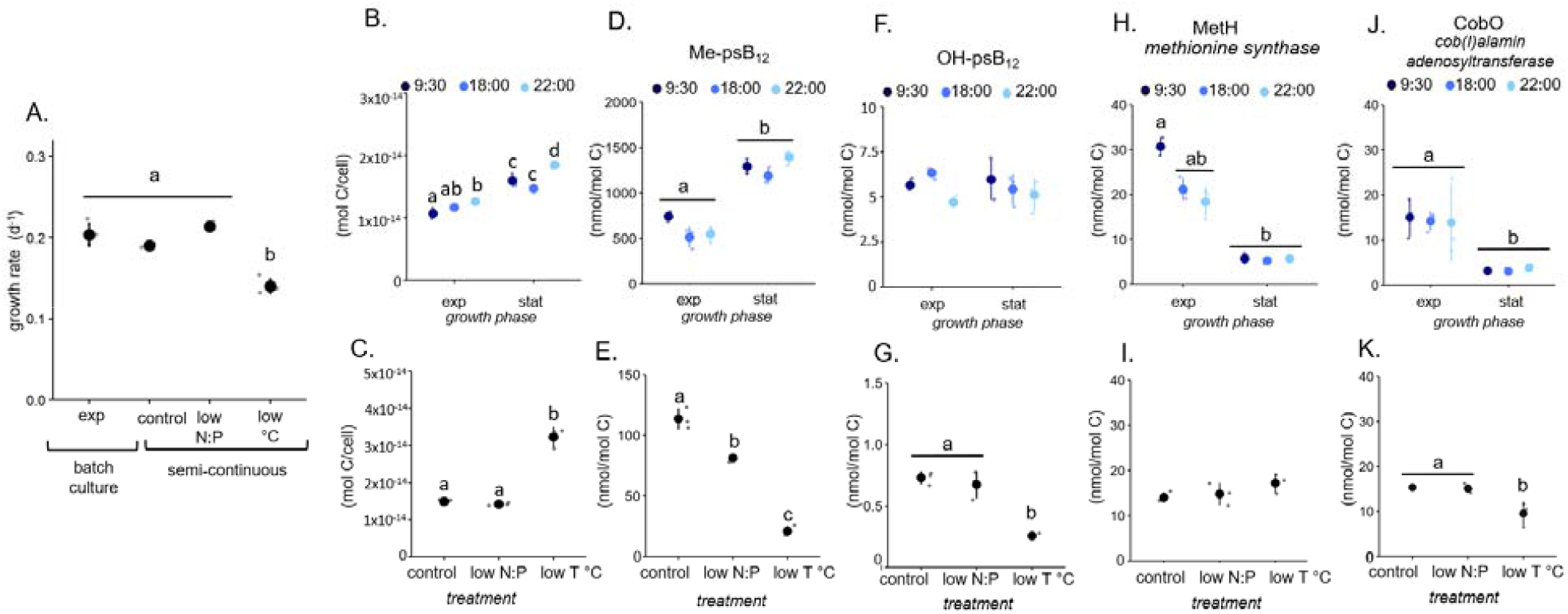
Changes in (A) growth rate (per day), (B,C) mol C per cell, (D, E) methyl-, (F, G) hydroxy- pseudocobalamin and (H,I) MetH and (J,K) CobO (nmol per mol carbon) in *Synechococcus* sp. WH8102 under different experimental treatments grown in batch (B, D, F, H, J) and in semi-continuous cultures (C, E, G, I, K). Mean values are represented by solid circles, individual biological replicate (n=3) values by small open circles, and the standard deviation by vertical lines. Different letters over data points indicate statistically significant differences (p-value < 0.05) between pairs of means based on the post-hoc Tukey’s Test. Shared letters (i.e. a, b, c) indicate no significant difference, if no letters are displayed then the differences between treatments were insignificant. *Synechococcus* sp. WH8102 culture data provided in Dataset 1.

Cellular Me-psB_12_ stoichiometry and quota (intracellular nutrient pool) ranged from 20 to 1500 nmol psB_12_ per mol C (Fig 2D, E) and 340 to 16,000 Me-psB_12_ molecules per cell respectively (Dataset 1). OH-psB_12_ ranged from 7 to <1 nmol psB_12_ per mol C in all samples (Fig 2F, G), 4 to 67 molecules per cell, and accounted for <1% of total psB_12_ inventory per cell (Fig 2 D-G). Despite our low limit of detection (LOD) (22 molecules per cell, Table S3), we did not detect Ado-psB_12_. Me-psB_12_ quota did not differ across diel cycle but was significantly higher in stationary phase (Fig 2D). Both Me-psB_12_ and OH-psB_12_ quotas increased >350% in cultures grown in batch compared to the control cultures grown semi-continuously although there was no difference in growth rate (Fig 2D-G). For semi-continuously grown cultures, Me-psB_12_ cellular quota significantly decreases at 17 °C compared to 24 °C, and when grown at 10 N:P versus the 26 N:P control (Fig 2E). In contrast, we found that OH-psB_12_ quota did not increase in stationary phase and was only significantly lower when grown semi-continuously at 17 °C (Fig 2F, G).

MetH peptide concentration ranged from 5 to 33 nmol per mol C in conditions tested (Fig 2H, I), representing 0.07 to 0.18% of total cellular protein (supplemental methods), and 40 to 370 MetH copies per cell (Dataset 1). MetH peptide concentration increased in the morning (9:30) at exponential phase when compared samples taken in the afternoon (18:00) and evening (22:00) (Fig 2H). MetH abundance (fmol per nmol carbon) did not vary in low temperature and low nitrogen (Fig 2I). CobO concentration ranged from 3 to 24 nmol per mol C in conditions tested (Fig 2J, KI), representing a 0.006 to 0.02% of total cellular protein (supplemental methods), and 20 to 240 CobO copies per cell (Dataset 1). CobO concentration was higher in exponential phase than stationary phase and were significantly lower in cultures grown in low temperature (Fig 2K). CobO fmol per µg protein was invariant in the conditions tested (Fig S3J, K).

### psB_12_ production and use by cyanobacteria in the Northwest Atlantic Ocean

To investigate psB_12_ production and use in the Northwest Atlantic Ocean, we measured cyanobacterial cells and particulate psB_12_ concentrations at four stations on the Scotian Shelf (HL 01, 02) and Slope (HL 05.5, 06) (Fig 3A). Peak chlorophyll *a* fluorescence (green) was between 10-30 m for HL 01 and HL 02 stations but were below 40 m for HL 05.5 and HL 06. Temperature decreased with depth at all stations but increased in surface waters from coastal to shelf break stations. Additional environmental data is provided in Dataset 2.

**Figure 3:**
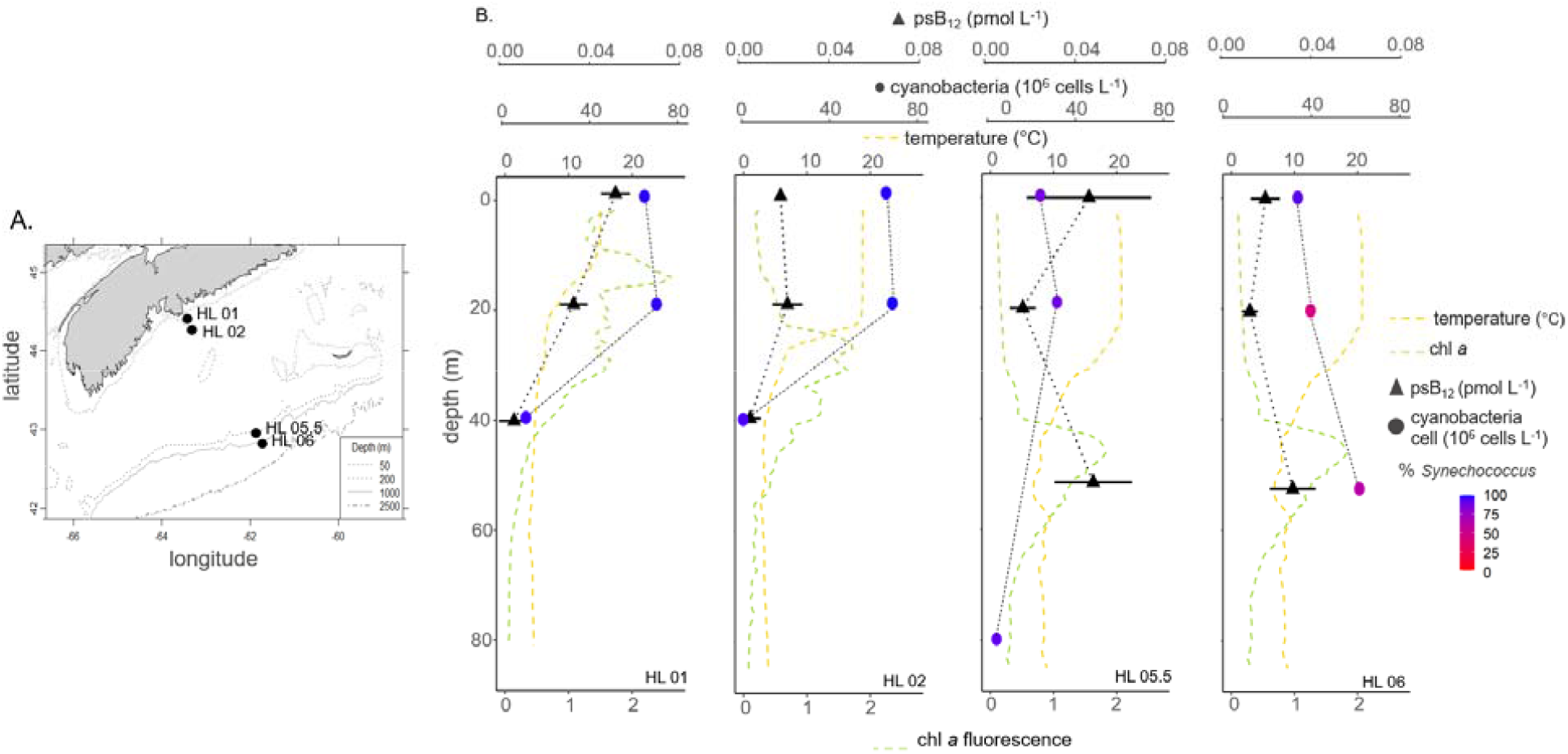
(A) Location of sampling on the Scotian Shelf and (B) depth profiles of particulate methyl-psB_12_ (pM, triangle), cyanobacterial abundance (10^6^ cells per L, circles), temperature (°C, yellow) and chlorophyll *a* fluorescence (green) depth profiles in the Fall, 2016. For pseudocobalamin, error bar represents technical replicates of mass spectrometry measurements. Circles are shaded based on the calculated % population of *Synechococcus* cells amongst all cyanobacteria (*Synechococcus* plus *Prochlorococcus*) determined through flow cytometry. Environmental data provided in Dataset 2.

Me-psB_12_ was the only quantifiable particulate psB_12_ form, despite our method’s low limits of detection for both OH- and Ado-psB_12_ (Table S3). Me-psB_12_ measurements were up to 0.08 pM and were undetectable below 50 m depth (Fig 3B). We found a higher particulate psB_12_ concentration at shallow depths and coastal waters, which generally mirrored cyanobacterial distribution. We also measured particulate B_12_ which ranged from undetectable to 0.4 pM total, with Me-B_12_, Ado-B_12_, and OH-B_12_ dominant in that order (Dataset 2).

*Synechococcus* cells per L were highest in the surface at HL 01 and HL 02. *Synechococcus* dominated cyanobacterial communities in most samples except for HL 06 at 20 m and 50 m (32% and 65% respectively) where *Prochlorococcus* also contributed (calculated from cyanobacteria cell counts, Dataset 2). *Synechococcus* comprised >65% of the cyanobacterial community in 1 m samples from all stations and these samples were therefore chosen to be screened for the *Synechococcus*-specific MetH peptide.

We examined the presence of the MetH peptide in environmental *Synechococcus* communities, leveraging a previous metagenomics study which identified potential prokaryotic cobalamin producers on the Scotian Shelf (Soto et al., 2023). Two *Synechococcus* genome bins (BB16_Bin_49-*Synechoccus* and AZOF_MAG_68-*Synechoccus*) are present at surface depths at HL 02 and HL 06 in Fall 2016 (Soto *et al*., 2023) (Fig 4A). The peptide we used to quantify MetH in *Synechococcus* in this study, highlighted in gray, is present in these bins (Fig 4A) and within available marine *Synechococcus* genomes from clade III, IV and I which are the dominant *Synechococcus* clades present in this region (Robicheau et al, 2022) (Fig 4B). The high contribution of cobalamin-associated reads from these two dominant *Synechococcus* bins at HL 02 and HL 06 in Fall 2016, and the presence of the MetH peptide in these bins, suggests that the MetH peptide we used in this study is likely to capture the majority of *Synechococcus* MetH present in these locations.

**Figure 4:**
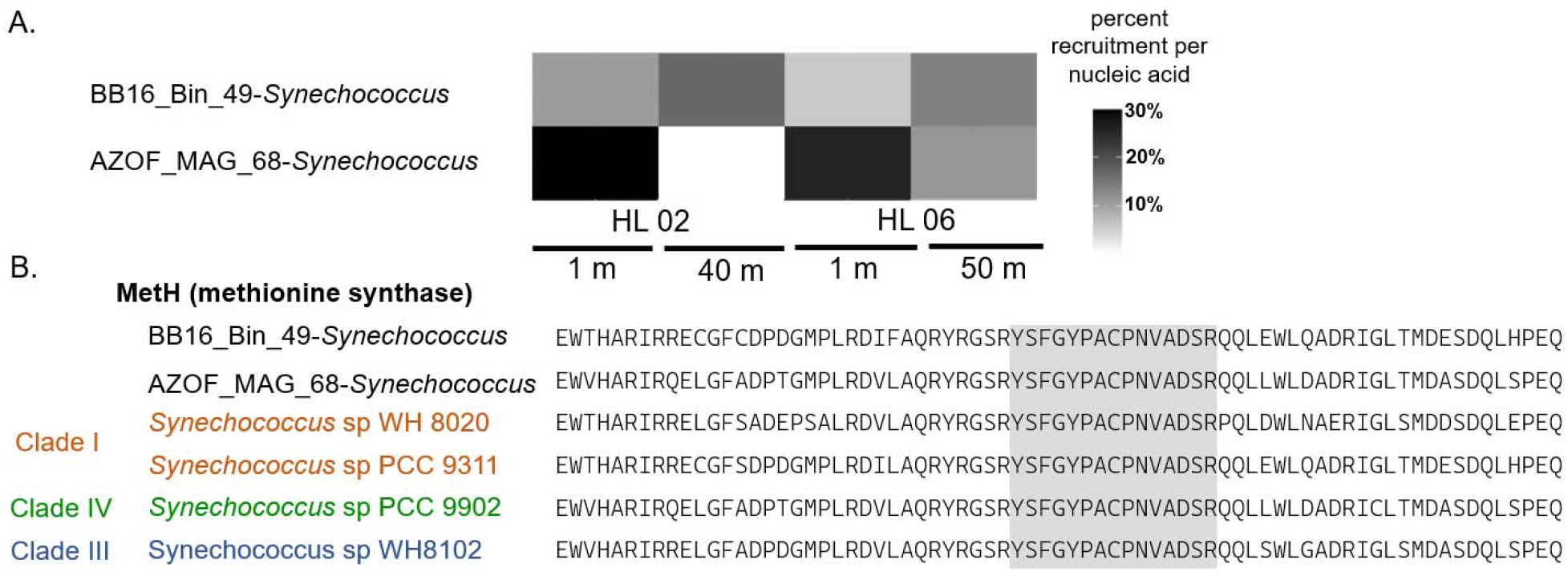
Assessing MetH peptide coverage in Scotian Shelf *Synechococcus* communities. (A) Recruitment of cobalamin-related reads per nucleic acid by two dominant *Synechococcus* metagenome assembled bins, shown as a proportion of all cobalamin-associated reads from the entire 0.2 um prokaryotic community at HL 02, HL 06 in Northwest Atlantic Ocean in Fall, 2016 (see Soto et al, 2023). (B) Alignment of MetH protein sequences from the two dominant assembled *Synechococcus* genome bins from Halifax Line in Fall 2016 (black) and representative *Synechococcus* genomes from Clade I (orange), Clade III (blue), Clade IV (green). Peptide used to quantify *Synechococcus* MetH in this study is highlighted in grey.

Assuming that all Me-psB_12_ originates from cyanobacteria, cyanobacteria in our samples had a Me-psB_12_ quota of 300 ± 100 molecules Me-psB_12_ per cell, with the exception of HL 05.5 at 1 m where there was ∼1150 Me-psB_12_ molecules per cyanobacterial cell (Fig 5A). Cyanobacterial cells count was used in place of *Synechococcus* cell counts as *Prochlorococcus* contributed to the cyanobacterial community in some of these samples (i.e. HL 06, Fig 3B). All *in situ* quotas fall into a range set by culture studies (Fig 2D, E). We also found an average of 460 ± 75 molecules of MetH per *Synechococcus* cell in surface samples measured (Fig 5B).

**Figure 5:**
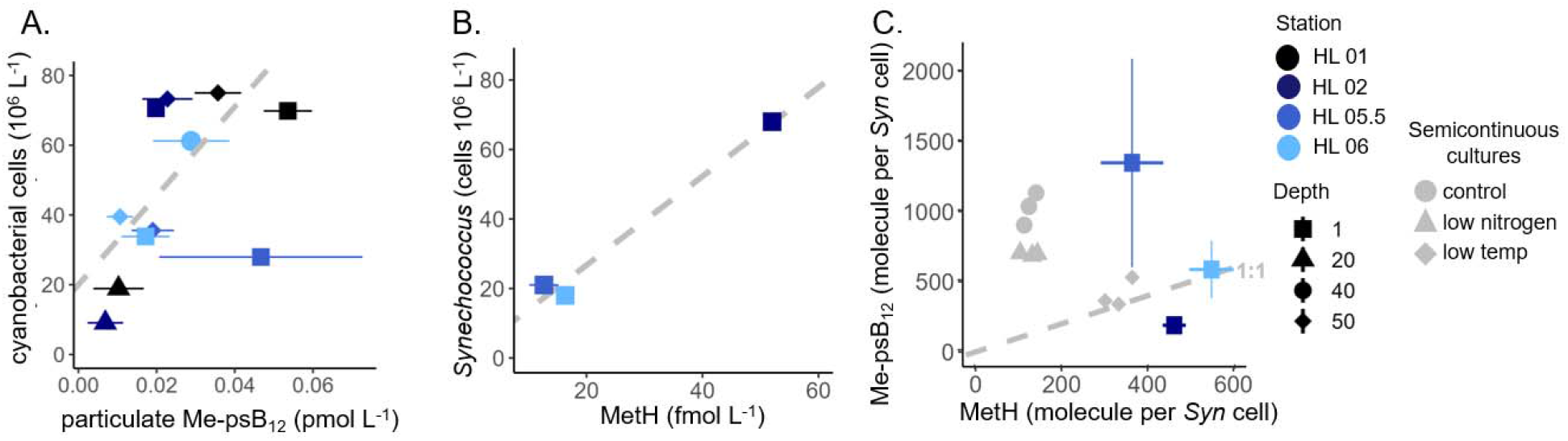
Pseudocobalamin production and use in the Northwest Atlantic Ocean. Relationship between (A) particulate psB_12_ (pM) and cyanobacterial cell counts (10^6^ L^-1^) in environmental samples. (B) Correlation between *Synechococcus* cell (10^6^ L^-1^) and *Synechococcus* MetH abundance (fmol L^-1^) in the surface ocean, (C) Protein (MetH) to cofactor (Me-psB_12_) ratio per *Synechococcus* cell in semicontinuous culture studies (grey circle, triangle, diamond) and environmental samples (blue squares). Horizontal and vertical error bars represent technical replicates measured using LC-MS/MS.

We then investigated the MetH:Me-psB_12_ stoichiometry in our culture and environmental community samples. Stoichiometry allows us to determine the proportion of Me-psB_12_ quota that is theoretically being used by MetH and provides insights into the *in situ* psB_12_ related status of the community and how it compared to cultures. Crystal structures of MetH have revealed that there is a 1:1 MetH to Me-B_12_ stoichiometry, and that MetH exclusively uses one Me-B_12_ molecule as a co-factor (Matthews *et al*., 2008; Koutmos *et al*., 2009). Although the structure has not been explicitly determined for cyanobacterial MetH, a study has demonstrated that cobalamin binding pocket of cyanobacterial psB_12_ accommodates the adenine of psB_12_ in place of the DMB (Heal *et al*., 2017). From this, we assume that MetH and psMe-B_12_ have a 1:1 protein to cofactor.

Notably, the sample from HL 02 was the only one to have a ratio lower than 1:1 (HL 02 = 0.4 Me-psB_12_: 1 MetH). This suggests that there was insufficient Me-psB_12_ to occupy the MetH enzymes present. In contrast, HL 05.5 had 2.8 Me-psB_12_:1 MetH. Both *Synechococcus* sp. WH8102 samples grown semi-continuously at 17 °C (low temp treatment) and environmental samples from HL 06 have close to the theoretical ratio of 1:1, indicating tight coupling of Me- psB_12_ production and use in these samples. Cultures grown in batch had significantly more Me- psB_12_ per cell than environmental samples and where therefore omitted from this graph but are displayed in Fig S4.

## Discussion

### Is protein abundance a suitable proxy for psB_12_ quotas?

There is increasing interest in leveraging transcriptomic and proteomic data to infer rates and processes in microbial communities (Hawco *et al*., 2020; Saito *et al*., 2020; Roberts *et al*., 2024). In theory, measuring coenzyme use (MetH) or production (CobO) could help approximate cellular concentrations or production rates of Me-psB_12_. In practice, this requires a thorough understanding of psB_12_ production, use, and changes in cofactor to protein ratios.

Here we show that CobO peptide per mol carbon did not always mirror psB_12_ quotas. For example, in stationary phase CobO was the lowest while Me-psB_12_ was the highest (Fig 2). Interestingly, CobO abundance and psB_12_ decreased in unison under low temperatures. However, CobO is just one step of approximately 30 that are in the pathway to produce corrinoids *de novo* (Warren *et al*., 2002, Fig 1B), and is responsible for the production of Ado- psB_12_ which we were unable to measure in both culture and environmental samples. Contrary to what can be observed in many heterotrophic bacteria, Cob/Cbi genes in *Synechococcus* (including the strain used in this study) are not clustered in operons and therefore may not be tightly co-regulated in *Synechococcus* (Rodionov *et al*., 2003). The same environmental or physiological condition may thus have a different effect on different Cob/Cbi genes. Recent research determined that CobSV, CobQ and CobW are key genes that enhanced cobalamin synthesis in *Ensifer adhaerens* HY-1 when more highly expressed (Xu *et al*., 2022). Future investigations into these genes, and others, may provide insight into the rate limiting step and regulation of psB_12_ and B_12_ biosynthesis.

MetH peptide abundance, perhaps a proxy for Me-psB_12_ use, rarely followed the general variations we observed in Me-psB_12_ in batch grown *Synechococcus* sp. WH8102 cultures. For example, in stationary phase MetH abundance was lowest while Me-psB_12_ was highest (Fig 2D, H). In semi-continuously grown cultures, MetH was invariant while Me-psB_12_ was significantly decreased in low N:P and temperature treatments (Fig 2E, I). However, the protein:co-factor stoichiometry for cultures grown at 17 °C were surprisingly close to 1:1, suggesting that under certain conditions inferring Me-psB_12_ quota from MetH abundance may be reasonable (Fig 5C). Although neither MetH nor CobO abundance sufficiently estimated or corresponded to Me-psB_12_ quota in cultured *Synechococcus* sp. WH8102, the observed protein and co-factor variability still provides insight into psB_12_-related cellular biology.

### Variability in psB_12_ quotas and protein inventories in Synechococcus sp. WH8102 cultures

Culture experiments here were performed on *Synechococcus* sp. WH8102 (clade III), a representative strain of a dominant clade in the region (Robicheau et al, 2022) to determine which variables may affect psB_12_ production and use. While not representative of all *Synechococcus* in this region, this culture work provides the first insights into the plasticity of psB_12_ quota and use in *Synechococcus* which can help contextualize environmental measurements. However, there is significant within and between clade variability in even cell size and macronutrient quotas (Harcourt et al, 2024), and so within and between-clade variability in psB_12_ is also expected. In general, further investigation into the relationship between different factors and psB_12_ related activity in different clades is an area of interest that can be pursued leveraging the peptides described here.

The relationship between temperature and psB_12_ quota is particularly important when conceptualizing the spatio-temporal patterns psB_12_ inventory and dynamics. Here, we demonstrate that *Synechococcus* sp. WH8102 grew significantly slower at 17 °C compared to the control 24 °C (Fig 2A), a result that is consistent with previous reports on this strain (Mackey *et al*., 2013; Varkey *et al*., 2016). *Synechococcus* sp. WH8102 cultures grown at low temperatures have a significant increase of mol carbon per cell (Fig 2B), possibly due to an increased cell size in lower temperatures (Moran et al, 2010) or decreased growth rate (Finkel et al, 2009), or both. Previous research has shown that C, N, and P quotas in various *Synechococcus* strains increase with decreased growth rate (Harcourt et al, 2024). In contrast, we show that mol Me- and OH-psB_12_ per mol carbon (Fig 2C-K) and molecules of psB_12_ per cell (Fig 5C, Dataset 2) significantly *decreased* in cultures grown at 17 °C compared to the control 24 °C. The decrease in psB_12_ per cell at lower temperatures suggests that there is 1) reduced requirements for psB_12_ or 2) a decrease in psB_12_ production, or both. In the low temperature culture, we did not observe a change in MetH but we did observe a significant decrease in CobO at low temperature (Fig 2H, J). This suggests that MetH abundance, and possibly psB_12_ demand, remains constant, and that instead perhaps enzymes involved in psB_12_ production may be more impacted by low temperature than most enzymes, creating a metabolic bottleneck in psB_12_ production. We speculate that this could be a result of high sensitivity to low temperature in the psB_12_ production pathway compared to other pathways. Or, potentially, during low temperature growth, resources could be allocated away from psB_12_ production. Given that the thermal optima for growth is variable across *Synechococcus* strains, further investigation into the relationship between growth rate, temperature, size, and psB_12_ quota in different clades is an area of interest.

We also demonstrate that psB_12_ quota in *Synechococcus* sp. WH8102 significantly decreased in low N:P, despite no change in growth rate or significant change in demand (MetH abundance) (Fig 2A, C). This result is of potential importance for the Scotian Shelf region, as a decrease in dissolved nitrogen relative to phosphorus has been observed since 2010 and is expected to continue in the future (Lehmann *et al*., 2023). On average, the N:P ratio in environmental samples was (3.5 ± 4.5 N:P) (Table S2) however no correlation between N:P and psB_12_ (pM) was observed. Further investigations of the relationships between N:P ratios, psB_12_ production, and remodeling activity are warranted, especially given that *B*_*12*_ and nitrogen have been shown to co-limit productivity and influence community composition in various oceanic regions, including the coast of the Gulf of Alaska (Koch *et al*., 2011), Long Island Embayment (Gobler *et al*., 2007) and Eastern boundary of the South Atlantic gyre (Browning *et al*., 2017)..

Me-psB_12_ (nmol per mol carbon) was significantly higher in *Synechococcus* sp. WH8102 harvested at stationary phase compared to exponential phase (Fig 2B). In stationary phase, there were 20 Me-psB_12_ for every MetH, suggesting that there are significant pools of psB_12_ in cells that are not associated with MetH (Dataset 1, Fig S4). This striking result points to either intracellular storage or alternative uses of Me-psB_12_ in *Synechococcus*, neither of which have been investigated in psB_12_ producers. Although roles for psB_12_ remain unknown, some alternative roles of *B*_*12*_ in the cell are beginning to be uncovered. For example, B_12_ is a sensitive light receptor and transcription regulator, albeit the benefit of these functions in stationary phase is not obvious (Ortiz-Guerrero *et al*., 2011; Cheng *et al*., 2016). Although increased psB_12_ quota in stationary phase was unexpected, this trend is similar to that of some harmful algal biotoxin producers where biotoxins accumulate in stationary phase for which reasons remain unclear (Tong *et al*., 2011; Song *et al*., 2023). Investigating the reservoirs of storage, accumulation, and alternative roles of psB_12_ could provide additional insight into the cellular processes that impact psB_12_ concentrations in cells and psB_12_ inventory in the ocean.

We observed a surprising >350% increase of psB_12_ quota in cultures grown in batches compared to cultures grown semicontinuously (Fig 2B, C; Fig 6A). Although psB_12_ quota at the beginning of the culture experiments was not measured, it could be expected that batch grown cells may have begun the experiment with a higher psB_12_ quota after accumulation in previous stationary phases (as seen in Fig 2D). Semi-continuously grown cultures were harvested after at least 7 generations (Fig S1) that did not reach stationary phase, and as such may have depleted their psB_12_ reserve. Unlike semi-continuous cultures, no new media was added to batch cultures during growth, therefore the higher psB_12_ concentrations may be due to the accumulation of either passively or actively released psB_12_ in the media or, potentially, the buildup of substances that may promote psB_12_ production. Unfortunately, we were unable to test this hypothesis due to the difficulties of measuring psB_12_ in the dissolved (<0.2 µm) phase.

**Figure 6:**
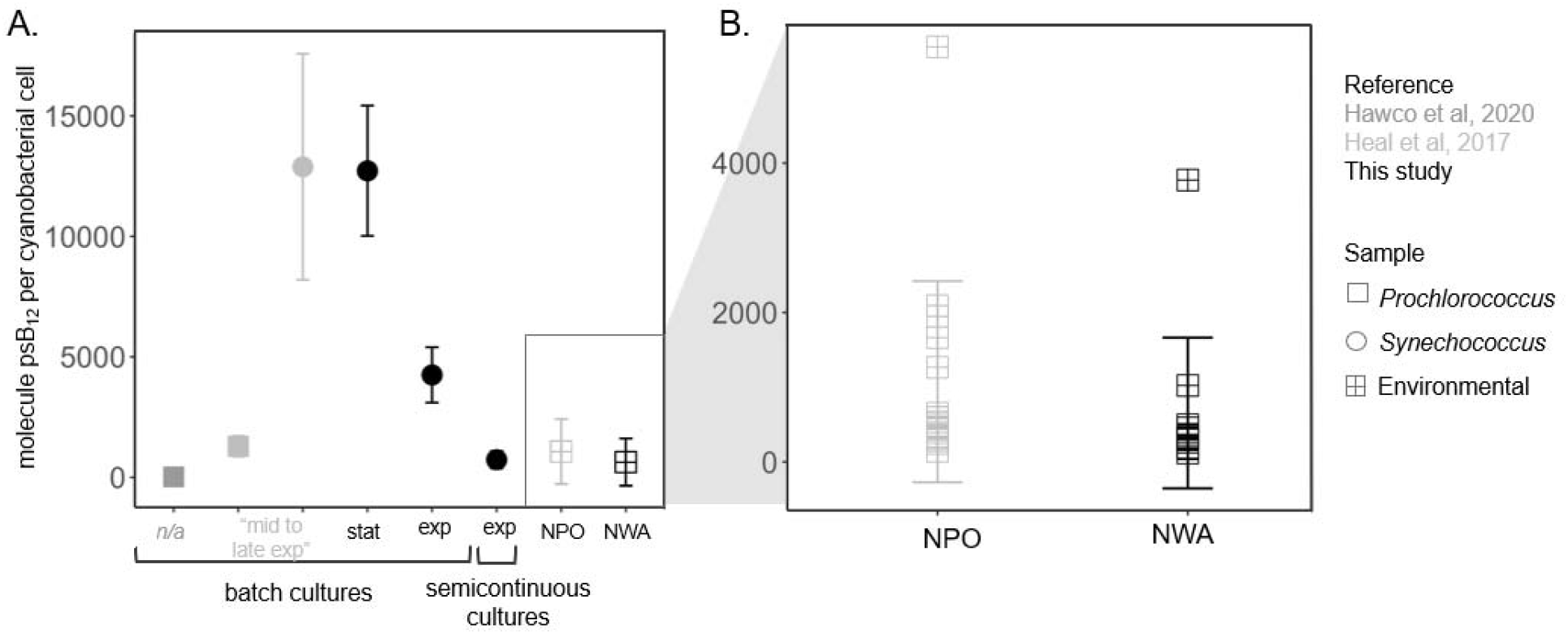
psB_12_ per cell in (A) cyanobacterial cultures and (B) the environment. (A) PsB_12_ per cell reported by Hawco et al, 2020* (dark gray), Heal et al, 2017 (light grey) and this study (black) in *Prochlorococcus* (square) and *Synechococcus* (circle) strains grown in batch or semicontinuously. Exp = exponential phase, stat = stationary phase. (B) psB_12_ quotas from Heal et al, 2017 from the North Pacific Ocean (NPO) (light grey) and from the Northwest Atlantic Ocean (NWA) (black) included in this study. *psB_12_ was inferred from targeted proteomic measurements of MetH

Interestingly, we observed the same influence of culture method when investigating *B*_*12*_ quota a recently isolated flavobacterium, *Pibocella* sp.. *Pibocella sp*. is a facultative B_12_ consumer, it does not make nor absolutely require B_12_, but accumulates significantly more B_12_ per cell when grown in batch compared to semi-continuously – even at the same dissolved B_12_ concentration (supplemental methods, Fig S5). Moreover, an increase in pg B_12_ per µg carbon was also observed when a haptophyte, *T. lueta*, was grown in batch cultures compared to in chemostats when grown at the same B_12_ concentration (Nef et al, 2019). The striking influence culture method has on both B_12_ and psB_12_ quota across different species merits further research and begs the question - *Which culture method best represents different environmental conditions?*

### Semi-continuous cultivation: most comparable to *in-situ* quotas

Comparing micronutrients quota between lab-grown cultures and environmental communities is rare but interest is growing. To determine which culture method of *Synechococcus* sp. WH8102 best reflected psB_12_ dynamics in an environmental context, we first compared psB_12_ quotas per *Synechococcus* cell. We found that psB_12_ per cell of batch grown cultures here fell in the same range of *Synechococcus* sp. WH8102 (clade III) and *Synechococcus* sp. WH7803 (clade V) reported in literature (Heal et al, 2017) (Fig 6A). However, batch cultures had much greater psB_12_ per cell concentrations than those measured semicontinuously grown cultures and in the environment (Fig 6B). We also compared the MetH to Me-psB_12_ stoichiometry from both culture and environmental studies. We found once again that samples from semi-continuously grown *Synechococcus* sp. WH8102 most closely resembled the *in-situ* stoichiometry of *Synechococcus* communities in this region (Fig 5C). Semi-continuously grown *Synechococcus* sp. WH8102 ranged from 1.2 to 8.2 Me-psB_12_ per MetH while environmental ranged from 0.4 to 2.8 Me-psB_12_ per MetH (Fig 5C). These results combined suggest that semi-continuously grown cultures appeared to be the most appropriate experimental approach for obtaining environmentally relevant data regarding psB_12_ in *Synechococcus* communities in our region (Fig 5C). Our results highlight that careful consideration is required when trying to infer metabolite dynamics in the field through lab cultures.

### Using cyanobacterial cell counts as a proxy for psB_12_ inventory

In this study, we provide the first set of environmental psB_12_ measurements in the Atlantic Ocean (Fig 3, 5). Here, psB_12_ accounted for 9 to 23% of total particulate cobalamin-related compounds (Fig7A, Table S2) - a significant part of total cobalamin that is missed if only B_12_ is measured. Missing psB_12_ measurements, especially in regions where cyanobacteria are abundant, restricts our ability to evaluate the importance psB_12_ and remodeling have on cobalamin production. However, measuring psB_12_, like many marine metabolites, is inherently challenging due to low concentration in a complex matrix, and lack of commercially available standard (Moran *et al*., 2022). Therefore, a proxy to estimate psB_12_ inventory could be a useful tool for researchers and further our understanding of when and how psB_12_ can influence marine microbial communities.

Here we found that 91% of our samples (10/11) contained 300 ± 100 Me-psB_12_ molecules per cyanobacteria cell (Fig 5A). The range of psB_12_ per cyanobacteria cell quantified in study was similar to the range of environmental measurements made from samples in the North Pacific Ocean (Heal *et al*., 2017) (Fig 6B). The consistency of psB_12_ measured inspired us to consider whether cyanobacterial cell counts may be sufficient to predict psB_12_ inventory. For this case study, we estimated total psB_12_ inventory over years at station HL 02 based *Synechococcus* cell counts from (Li, 2014), prepared as described in (Li and Dickie, 2001) (Fig 7B). While we used a narrower quota for this case study, we suggest that a general order of 10^3^ psB_12_ molecules per cyanobacterial cell could be useful for estimating psB_12_ inventory from cyanobacterial cell counts.

**Figure 7:**
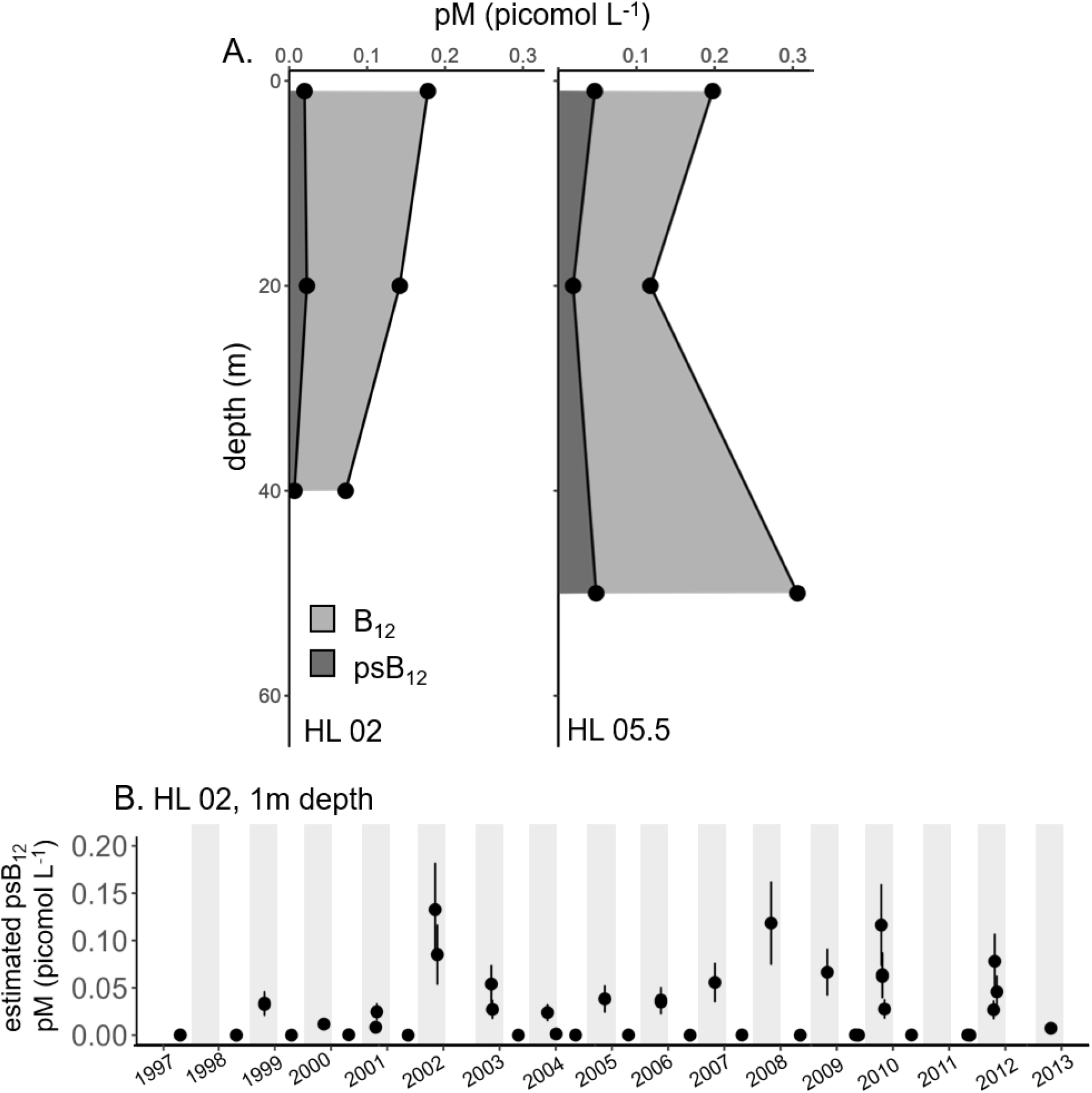
(A) psB_12_ (dark gray) and combined (light gray) total particulate B_12_ (pM) down the water column at HL 02 and HL 05.5. Environmental data presented in Table S2. (B) Estimated particulate psB_12_ inventory (pM) from 1997-2013 at HL 02 (1m) calculated from S*ynechococcus* cell counts from Li, 2014.

The 10^3^ psB_12_ molecules per cyanobacterial cell estimate suggested here is *a* quota, not *the* quota. Outliers from this range will undoubtedly appear with additional environmental measurements. However, just like the outliers from HL 05.5, we believe that these outliers provide valuable information that can guide future research. The value of psB_12_ per cyanobacterial cell at 1 m from HL 05.5 still fall into a reasonable range reported by this study and literature (Fig 6A, B) (Hawco et al, 2020; Heal et al, 2017). HL 05.5 is located on the Scotian Shelf break, an oceanographically complex region where the microbial community present is more diverse than other stations (Zorz *et al*., 2019). Indeed, when examining the previously analyzed cyanobacterial (16S) mapped ASVs of the samples used in this study (Fig S6), we see that samples from HL 05.5 had highest Shannon diversity and chao values compared to the others sampled (Fig S7) (Robicheau *et al*., 2022). This indicates that community diversity may contribute to the increased Me-psB_12_ cell quota seen in these samples – emphasizing the importance of psB_12_ measurements from a range of cyanobacteria and *Synechococcus* clades if more precise quota predictions are of utility.

## Conclusion

Although psB_12_ is a potentially significant source of B_12_ in the ocean, there are limited measurements of it, and the factors that influence its production and use are unknown. Here, we reveal that in culture conditions, psB_12_ quota decreases in low temperature and low N:P and that both growth phase and culturing methods can impact psB_12_ quota in *Synechococcus* sp. WH8102. In samples from the Northwest Atlantic Ocean, we found that psB_12_ quota per cyanobacteria cell was surprisingly consistent. This provides researchers with a proxy to systematically incorporate cyanobacteria and remodellers into conceptual and numerical models of environmental B_12_ dynamics by estimating psB_12_ inventory. These quantitative values are a step towards understanding of the role psB_12_ has in marine microbial communities.

## Acknowledgements

Thanks to Dr. Jenni Tolman for the advice on processing flow cytometry samples and to Dr. William Li for the use of his long-term record of *Synechococcus* cell counts and his seminal work on picophytoplankton in this region. We are grateful to the Bedford Institute of Oceanography, Fisheries and Oceans Canada Atlantic Zone Monitoring Program (AZMP) for access to the sea for the collection of samples analyzed in this study. This study was funded by NSERC Discovery Grant RGPIN2015-05009 to EMB, Simons Foundation Grant 504183 to EMB, Simons Foundation CBIOMES Award ID 1001702 to EMB, Canada Research Chair Support to EMB, Ocean Frontier Institute support to EMB and JLR, and NSERC CGS-D to CB.

## References

Bertrand, E.M., McCrow, J.P., Moustafa, A., Zheng, H., McQuaid, J.B., Delmont, T.O., et al. (2015) Phytoplankton–bacterial interactions mediate micronutrient colimitation at the coastal Antarctic sea ice edge. Proc Natl Acad Sci USA 112: 9938–9943.

Browning, T.J., Achterberg, E.P., Rapp, I., Engel, A., Bertrand, E.M., Tagliabue, A., and Moore, C.M. (2017) Nutrient co-limitation at the boundary of an oceanic gyre. Nature 551: 242–246.

Cheng, Z., Yamamoto, H., and Bauer, C.E. (2016) Cobalamin’s (Vitamin B12) Surprising Function as a Photoreceptor. Trends Biochem Sci 41: 647–650.

Dupont, C.L., Barbeau, K., and Palenik, B. (2008) Ni Uptake and Limitation in Marine Synechococcus Strains. Appl Environ Microbiol 74: 23–31.

Flombaum, P., Gallegos, J.L., Gordillo, R.A., Rincon, J., Zabala, L.L., Jiao, N., et al. (2013) Present and future global distributions of the marine Cyanobacteria Prochlorococcus and Synechococcus. Proceedings of the National Academy of Sciences 110: 9824–9829.

Gleason, F.K. and Olszewski, N.E. (2002) Isolation of the Gene for the B12-Dependent Ribonucleotide Reductase from Anabaena sp. Strain PCC 7120 and Expression in Escherichia coli. Journal of Bacteriology 184: 6544–6550.

Gobler, C., Norman, C., Panzeca, C., Taylor, G., and Sañudo-Wilhelmy, S. (2007) Effect of B-vitamins (B1, B12) and inorganic nutrients on algal bloom dynamics in a coastal ecosystem. Aquat Microb Ecol 49: 181–194.

Gurdeep Singh, R., Tanca, A., Palomba, A., Van der Jeugt, F., Verschaffelt, P., Uzzau, S., et al. (2019) Unipept 4.0: Functional Analysis of Metaproteome Data. J Proteome Res 18: 606–615.

Hawco, N.J., McIlvin, M.M., Bundy, R.M., Tagliabue, A., Goepfert, T.J., Moran, D.M., et al. (2020) Minimal cobalt metabolism in the marine cyanobacterium Prochlorococcus. Proc Natl Acad Sci USA 117: 15740–15747.

Heal, K.R., Qin, W., Ribalet, F., Bertagnolli, A.D., Coyote-Maestas, W., Hmelo, L.R., et al. (2017) Two distinct pools of B <sub>12 </sub>analogs reveal community interdependencies in the ocean. Proc Natl Acad Sci USA 114: 364–369.

Helliwell, K.E., Lawrence, A.D., Holzer, A., Kudahl, U.J., Sasso, S., Kräutler, B., et al. (2016) Cyanobacteria and Eukaryotic Algae Use Different Chemical Variants of Vitamin B12. Current Biology 26: 999– 1008.

Hoofnagle, A.N., Whiteaker, J.R., Carr, S.A., Kuhn, E., Liu, T., Massoni, S.A., et al. (2016) Recommendations for the Generation, Quantification, Storage, and Handling of Peptides Used for Mass Spectrometry–Based Assays. Clinical Chemistry 62: 48–69.

Kido Soule, M.C., Longnecker, K., Johnson, W.M., and Kujawinski, E.B. (2015) Environmental metabolomics: Analytical strategies. Marine Chemistry 177: 374–387.

Koch, F., Marcoval, M.A., Panzeca, C., Bruland, K.W., Sañudo-Wilhelmy, S.A., and Gobler, C.J. (2011) The effect of vitamin B12 on phytoplankton growth and community structure in the Gulf of Alaska. Limnology and Oceanography 56: 1023–1034.

Koutmos, M., Datta, S., Pattridge, K.A., Smith, J.L., and Matthews, R.G. (2009) Insights into the reactivation of cobalamin-dependent methionine synthase. Proceedings of the National Academy of Sciences 106: 18527–18532.

Lehmann, N., Reed, D.C., Buchwald, C., Lavoie, D., Yeats, P.A., Mei, Z.-P., et al. (2023) Decadal Variability in Subsurface Nutrient Availability on the Scotian Shelf Reflects Changes in the Northwest Atlantic Ocean. Journal of Geophysical Research: Oceans 128: e2023JC019928.

Li, W.K.W. (2014) The state of phytoplankton and bacterioplankton on the Scotian Shelf and Slope: Atlantic Zone Off-Shelf Monitoring Program 1994-2013, Dartmouth, Nova Scotia: Can. Tech. Rep. Hydrogr. Ocean. Sci.

Li, W.K.W. and Dickie, P.M. (2001) Monitoring phytoplankton, bacterioplankton, and virioplankton in a coastal inlet (Bedford Basin) by flow cytometry. Cytometry 44: 236–246.

Li, W.K.W. and Wood, A.M. (1988) Vertical distribution of North Atlantic ultraphytoplankton: analysis by flow cytometry and epifluorescence microscopy. Deep Sea Research Part A Oceanographic Research Papers 35: 1615–1638.

Loder, J., Han, G., Galbraith, P., Chasse, J., and van der Baaren, A. eds. (2013) Aspects of climate change in the Northwest Atlantic off Canada. Canadian Technical Report of Fisheries and Aquatic Sciences 3045 202.

Lu, X., Heal, K.R., Ingalls, A.E., Doxey, A.C., and Neufeld, J.D. (2020) Metagenomic and chemical characterization of soil cobalamin production. ISME J 14: 53–66.

Lundin, D., Gribaldo, S., Torrents, E., Sjöberg, B.-M., and Poole, A.M. (2010) Ribonucleotide reduction - horizontal transfer of a required function spans all three domains. BMC Evol Biol 10: 383.

Ma, A.T., Tyrell, B., and Beld, J. (2020) Specificity of cobamide remodeling, uptake and utilization in Vibrio cholerae. Molecular Microbiology 113: 89–102.

Mackey, K.R.M., Paytan, A., Caldeira, K., Grossman, A.R., Moran, D., McIlvin, M., and Saito, M.A. (2013) Effect of Temperature on Photosynthesis and Growth in Marine Synechococcus spp. Plant Physiology 163: 815–829.

Mackey, K.R.M., Post, A.F., McIlvin, M.R., Cutter, G.A., John, S.G., and Saito, M.A. (2015) Divergent responses of Atlantic coastal and oceanic Synechococcus to iron limitation. Proc Natl Acad Sci U S A 112: 9944–9949.

Matthews, R.G., Koutmos, M., and Datta, S. (2008) Cobalamin-dependent and cobamide-dependent methyltransferases. Current Opinion in Structural Biology 18: 658–666.

Moran, M.A., Kujawinski, E.B., Schroer, W.F., Amin, S.A., Bates, N.R., Bertrand, E.M., et al. (2022) Microbial metabolites in the marine carbon cycle. Nature Microbiology 7: 508–523.

Ortiz-Guerrero, J.M., Polanco, M.C., Murillo, F.J., Padmanabhan, S., and Elías-Arnanz, M. (2011) Light-dependent gene regulation by a coenzyme B <sub>12 </sub>-based photoreceptor. Proc Natl Acad Sci USA 108: 7565–7570.

Papadopoulos, J.S. and Agarwala, R. (2007) COBALT: constraint-based alignment tool for multiple protein sequences. Bioinformatics 23: 1073–1079.

Roberts, M.E., Bhatia, M.P., Rowland, E., White, P.L., Waterman, S., Cavaco, M.A., et al. (2024) Rubisco in high Arctic tidewater glacier-marine systems: A new window into phytoplankton dynamics. Limnology and Oceanography n/a:

Robicheau, B.M., Tolman, J., Bertrand, E.M., and LaRoche, J. (2022) Highly-resolved interannual phytoplankton community dynamics of the coastal Northwest Atlantic. ISME COMMUN 2: 1–12.

Rodionov, D.A., Vitreschak, A.G., Mironov, A.A., and Gelfand, M.S. (2003) Comparative Genomics of the Vitamin B <sub>12 </sub>Metabolism and Regulation in Prokaryotes. J Biol Chem 278: 41148–41159.

Saito, M.A., McIlvin, M.R., Moran, D.M., Santoro, A.E., Dupont, C.L., Rafter, P.A., et al. (2020) Abundant nitrite-oxidizing metalloenzymes in the mesopelagic zone of the tropical Pacific Ocean. Nat Geosci 13: 355–362.

Song, W., Song, X., Chi, L., Zhu, J., Cao, X., and Yu, Z. (2023) Novel insights into toxin changes associated with the growth of Alexandrium pacificum: Revealing active toxin-secretion ability and toxin cell quota variation. Harmful Algae 129: 102516.

Soto, M.A., Desai, D., Bannon, C., LaRoche, J., and Bertrand, E.M. (2023) Cobalamin producers and prokaryotic consumers in the Northwest Atlantic. Environmental Microbiology n/a:

Suffridge, C.P., Gómez-Consarnau, L., Monteverde, D.R., Cutter, L., Arístegui, J., Alvarez-Salgado, X.A., et al. (2018) B Vitamins and Their Congeners as Potential Drivers of Microbial Community Composition in an Oligotrophic Marine Ecosystem. Journal of Geophysical Research: Biogeosciences 123: 2890–2907.

Tanioka, Y., Miyamoto, E., Yabuta, Y., Ohnishi, K., Fujita, T., Yamaji, R., et al. (2010) Methyladeninylcobamide functions as the cofactor of methionine synthase in a Cyanobacterium, Spirulina platensis NIES-39. FEBS Letters 584: 3223–3226.

Tanioka, Y., Yabuta, Y., Yamaji, R., Shigeoka, S., Nakano, Y., Watanabe, F., and Inui, H. (2009) Occurrence of Pseudovitamin B12 and Its Possible Function as the Cofactor of Cobalamin-Dependent Methionine Synthase in a Cyanobacterium Synechocystis sp. PCC6803. J Nutr Sci Vitaminol 55: 518–521.

Tong, M., Kulis, D.M., Fux, E., Smith, J.L., Hess, P., Zhou, Q., and Anderson, D.M. (2011) The effects of growth phase and light intensity on toxin production by Dinophysis acuminata from the northeastern United States. Harmful Algae 10: 254–264.

Varkey, D., Mazard, S., Ostrowski, M., Tetu, S.G., Haynes, P., and Paulsen, I.T. (2016) Effects of low temperature on tropical and temperate isolates of marine Synechococcus. ISME J 10: 1252– 1263.

Warren, M.J., Raux, E., Schubert, H.L., and Escalante-Semerena, J.C. (2002) The biosynthesis of adenosylcobalamin (vitamin B12). Nat Prod Rep 19: 390–412.

Xu, S., Xiao, Z., Yu, S., Zeng, W., Zhu, Y., and Zhou, J. (2022) Enhanced cobalamin biosynthesis in Ensifer adhaerens by regulation of key genes with gradient promoters. Synthetic and Systems Biotechnology 7: 941–948.

Zorz, J., Willis, C., Comeau, A.M., Langille, M.G.I., Johnson, C.L., Li, W.K.W., and LaRoche, J. (2019) Drivers of Regional Bacterial Community Structure and Diversity in the Northwest Atlantic Ocean. Front Microbiol 10: 281.

