## Supplement Information for "Pseudocobalamin production and use in marine *Synechococcus* cultures and communities"

**Document Includes:
*Supporting methods***

- Calculating proportion of total cellular protein in culture studies
- Mass spectrometry analysis details
- *Pibocella* sp. culturing

**Tables S1-4**
Table S1: Transition list for targeted mass spectrometry (proteomics)
Table S2: Transition list for targeted mass spectrometry (metabolomics)
Table S3: Limit of detection and quantification for targeted metabolomics

**Figures S1-S7**
Figure S1: Growth curves of *Synechococcus* sp. WH 8102 for culture studies
Figure S2: Flow cytogram of HL_06 at 20 m depth.
Figure S3: Changes in growth rate (per day), µg total protein per cell, Me-, OH- pseudocobalamin, and MetH and CobO *(fmol per µgram total protein)*
Figure S4: Full version of Fig 5C, MetH to Me-pseudocobalamin ratio in *Synechococcus* sp. WH 8102 culture study and environmental samples
Figure S5: Cobalamin quota of *Pibocella* isolate grown in cobalamin replete low nutrient heterotrophic media
Figure S6: Relative eukaryotic and cyanobacteria and relative cyanobacterial ASVs at HL01, HL02, HL05.5 and HL06 in large (>3 µm) and small (<3 µm, >0.22 µm) size fractions
Figure S7: Diversity indices for environmental samples

***Supplemental Methods***

***Calculations of percent of total cellular protein***

$$\frac{fmol MetH}{1 \mu g protein} x 132 100 g per mol (MW {MetH}^{*})=pg MetH per \mu g protein$$

$$\frac{pg MetH per \mu g protein}{{10}^{6}}= \mu g MetH per \mu g protein$$

$$\mu g MetH per \mu g protein x \mu g protein per filter= \mu g MetH per filter$$

$$\frac{\mu g MetH per filter}{\mu g protein per filter}=\% MetH of total protein$$

*equivalent in kDa

MetH
(methionine synthase [Synechococcus sp. WH 8102]), NCBI number WP_011128102.1 = 132.1 kDa

CobO
(MULTISPECIES: cob(I)alamin adenosyltransferase [Synechococcus]), NCBI number WP_011127151.1 = 20.9 kDa

***Mass spectrometry analysis details***
*Metabolomic MS set up*
 Conditions for metabolomic selective reaction monitoring were as followed: Q1 and Q3 resolution 0.7 (FWHM), 50 ms dwell time, spray voltage 3500 (positive ion mode), sheath gas 6, auxiliary gas 2, ion transfer tube 325 ºC, vaporizer temperature 100 ºC. Triplicate 5 µL injections were performed onto a 150 x 0.3 mm ID column (Acclaim PepMap RSLC, C-18, 2 µm, 100 Å) with a 5 x 0.3 mm ID guard column in front, held at 50 ºC and subject to an HPLC gradient of 2 – 32% B over 6 min, then 32 - 60% B over 0.5 min (A, 20 mM ammonium formate, 0.1% formic acid; B, 0.1% formic acid in acetonitrile) at 10 µl per min.
*Proteomic MS set up*
 The MS was operated with a spray voltage of 3500 V, sheath gas 5, auxiliary gas 2, ion transfer tube 325 ºC, vaporizer gas 70 ºC and a Chrom filter setting of 10 s. 1 µg protein samples were spiked with 20 fmol of each heavy isotope-labeled peptide standard and loaded onto 5 mm x 0.3 mm I.D. C18 trapping column at 20 µl/min and then separated over a 150 x 0.3 mm ID reverse phase column (Acclaim C18, 2 µm, 100 Å), 4 – 43% B over 40 min, 5 µl/min, 50 ºC. Mobile phase A was 0.1% formic acid; B was 80% acetonitrile, 0.08% formic acid. The method contained 80 transitions, 15 msec dwell time, Q1 and Q3 resolution was set to 0.7 (FWHM), automatically calibrated RF lens setting and a collision gas pressure 2.5 mTorr.

***Methods for culturing Pibocella sp. isolate***

For batch cultures, *Pibocella* sp. were grown in the dark at room temperature in low nutrient heterotrophic media (LNHM) (Davis and Guillard 1958; Cho and Giovannoni 2003) in quadruplicates and transferred every 6 days with a 1:100 dilution for a total of 2 transfers resulting in 23 generations. Cultures were grown semicontinuously in triplicates and transferred using a 1:100 dilution every second day for a total of 11 transfers and 46 generations (inferred from growth rate). Cell counts samples were collected every two days fixing 0.2 mL culture aliquots with 1% paraformaldehyde and storing at -20° C every two days during growth. Batch cultured cells were harvested in stationary phase and semi-continuously cultured cells were harvested in late exponential phase through filtration on a 0.22 µm nylon filter for metabolite extraction and frozen at -80 °C until extracted as described in manuscript. Before flow cytometry analysis 2 µL of 1:100 SYBR Green was added to fixed culture aliquots and incubated to stain bacterial cells. Cell counts for batch culture samples were performed on a BD Accuri™ C6 Flow Cytometer (BD Biosciences, San Jose, CA) and for semicontinuous culture samples cell counts were performed on a NovoCyte flow cytometer (Agilent Technologies, San Diego, CA). Growth rates (d^-1^) were derived from these cell counts and determined to be 0.65 ± 0.1 (batch) and 1.33 ± 0.02 (semicontinuous). Carbon content was calculated based on the partial density of carbon of 111.9 fg C µm^-3^ for marine bacteria (Arandia-Gorostidi et al. 2020; Fukuda et al. 1998) and the biomass of *Pibocella sp*. (Nedashkovskaya et al., 2005) resulting in an estimate of 32 fg carbon cell^-1^ falling within the lower and upper estimates of marine flavobacteria carbon content (Thomas et al., 2021).

***Supplemental Tables***

**Table S1:** Selected reaction monitoring mass spectrometry parameters for peptides measured in this study.

| **Peptide** | **Protein** | **Precursor (m/z)** | **Product (m/z)** | **Collision Energy (eV)** | **Retention time (min)** |
| --- | --- | --- | --- | --- | --- |
| YSFGYPAC[+57]PNVADSR(+2) | MetH | 852.4 | 1086.5 | 26.7 | 22.4 |
| YSFGYPAC[+57]PNVADSR(+2) |  | 852.4 | 918.4 | 28.4 | 22.4 |
| YSFGYPAC[+57]PNVADSR(+2) |  | 852.4 | 758.4 | 28.5 | 22.4 |
| YSFGYPAC[+57]PNVADSR(+2) |  | 852.4 | 618.3 | 22.6 | 22.4 |
| YSFGYPAC[+57]PNVADSR (heavy)(+2) |  | 855.4 | 1092.5 | 26.7 | 22.4 |
| YSFGYPAC[+57]PNVADSR (heavy)(+2) |  | 855.4 | 924.4 | 28.4 | 22.4 |
| YSFGYPAC[+57]PNVADSR (heavy)(+2) |  | 855.4 | 764.4 | 28.5 | 22.4 |
| YSFGYPAC[+57PNVADSR (heavy)(+2) |  | 855.4 | 618.3 | 22.6 | 22.4 |
| GLVLVFTGQGK(+2) | CobO | 559.8 | 849.5 | 17.8 | 25.4 |
| GLVLVFTGQGK(+2) |  | 559.8 | 736.4 | 18.3 | 25.4 |
| GLVLVFTGQGK(+2) |  | 559.8 | 637.3 | 17.5 | 25.4 |
| GLVLVFTGQGK(+2) |  | 559.8 | 389.2 | 17.2 | 25.4 |
| GLVLVFTGQGK (heavy)(+2) |  | 562.8 | 855.5 | 17.8 | 25.4 |
| GLVLVFTGQGK (heavy)(+2) |  | 562.8 | 742.4 | 18.3 | 25.4 |
| GLVLVFTGQGK (heavy)(+2) |  | 562.8 | 637.3 | 17.5 | 25.4 |
| GLVLVFTGQGK (heavy)(+2) |  | 562.8 | 389.2 | 17.2 | 25.4 |

**Table S2:** Selected reaction monitoring mass spectrometry parameters for metabolites measured in this study.

| **Cobalamin** | **Precursor (m/z)** | **Product (m/z)** | **Collision Energy (eV)** | **Retention time (min)** |
| --- | --- | --- | --- | --- |
| OH-psB_12_ | 659.8 | 136.1 | 30.0 | 4.1 |
|  | 659.8 | 348.1 | 30.0 | 4.1 |
| OH-B_12_ | 665.0 | 147.1 | 37.4 | 4.4 |
|  | 665.0 | 359.1 | 24.4 | 4.4 |
| Me-psB_12_ | 668.3 | 136.1 | 30.0 | 5.7 |
|  | 668.3 | 348.1 | 30.0 | 5.7 |
| Me-B_12_ | 673.5 | 147.1 | 40.8 | 6.1 |
|  | 673.5 | 359.1 | 26.0 | 6.1 |
| Ado-psB_12_ | 785.3 | 136.1 | 30.0 | 4.7 |
|  | 785.3 | 348.1 | 30.0 | 4.7 |
| Ado- B_12_ | 790.9 | 147.1 | 43.3 | 5.5 |
|  | 790.9 | 359.1 | 29.7 | 5.5 |
| CN-B_12_-heavy | 681.9 | 154.1 | 38.0 | 4.9 |
|  | 681.9 | 366.1 | 23.7 | 4.9 |
|  | 681.9 | 997.5 | 23.0 | 4.9 |
|  | 681.9 | 1210.5 | 22.7 | 4.9 |
|  | 681.9 | 639.4 | 21.2 | 4.9 |
|  | 681.9 | 912.5 | 35.0 | 4.9 |

**Table S3:** Limit of detection (LOD) and quantification (LOQ) for metabolites quantified in this study presented in fmol on HPLC column and molecule per cell (*Synechococcus sp*. WH 8102 samples) and pM (environmental samples)

| **psB_12_** | **B_12_ standard** | **LOD  (fmol on HPLC column)** | **LOQ  (fmol on HPLC column)** | **LOD  8102 = molecule per cell, environmental = pM** | **LOQ  8102 = molecule per cell, environmental = pM** |
| --- | --- | --- | --- | --- | --- |
| **8102 samples** | | | | | |
| Me-psB_12_ | Me-B_12_ | 0.59 | 1.98 | 12 | 40 |
| OH-psB_12_ | OH-B_12_ | 0.22 | 0.74 | 4 | 15 |
| Ado-psB_12_ | Ado-B_12_ | 1.09 | 3.66 | 22 | 73 |
| **Environmental samples** | | | | | |
| Me-psB_12_ | Me-B_12_ | 0.35 | 1.16 | 0.007 | 0.023 |
| OH-psB_12_ | OH-B_12_ | 0.35 | 1.18 | 0.007 | 0.024 |
| Ado-psB_12_ | Ado-B_12_ | 1.82 | 6.08 | 0.036 | 0.122 |

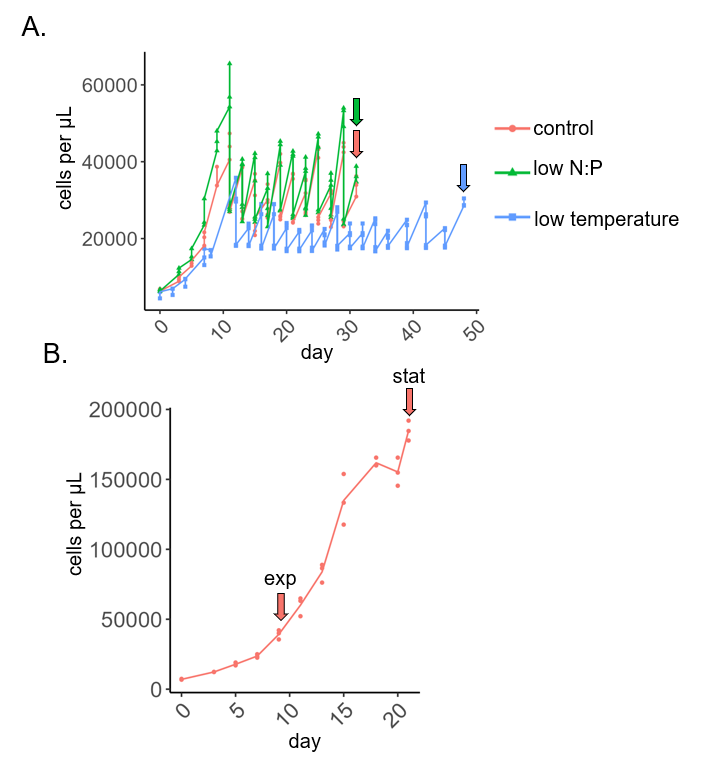

**Figure S1:** Growth curves of *Synechococcus* sp. WH 8102 cultures used in the study grown (A) semicontinuously and (B) in batch. Culture conditions are listed in *Table 1* and harvest dates are indicated by arrow. exp = exponential, stat = stationary.

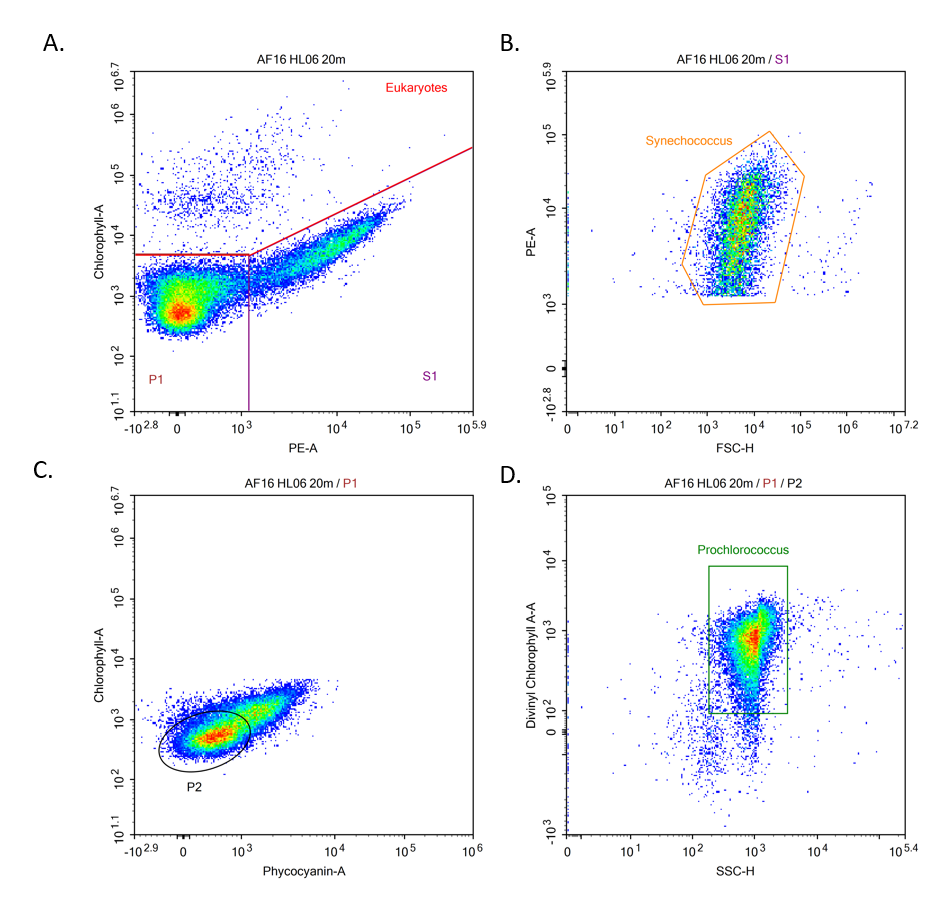
 **Figure S2:** Representative flow cytogram of sample from HL_06 at 20 m in Fall of 2016 A) chlorophyll-A (blue laser (488 nm); 675/30 m filter) and phycoerythrin (PE) (blue laser (488 nm);572/X nm filter) with P1 and S1 groups. B) Daughter plot of S1 in (A) with phycoerythrin (PE) (blue laser (488 nm);572/X nm filter) and forward scatter indicating *Synechococcus* gate used for counts. C) Daughter plot of P1 in (A) with chlorophyll-A (blue laser (488 nm); 675/30 m filter) and phycocyanin-A (red laser (640 nm); 675/30 nm filter) indicating P2. D) Daughter plot of P2 with divinyl chlorophyll-A (DChlA) (violet laser (405 nm); 675/30 nm filter) and side scatter indicating *Prochlorococcus* gate used for counts.

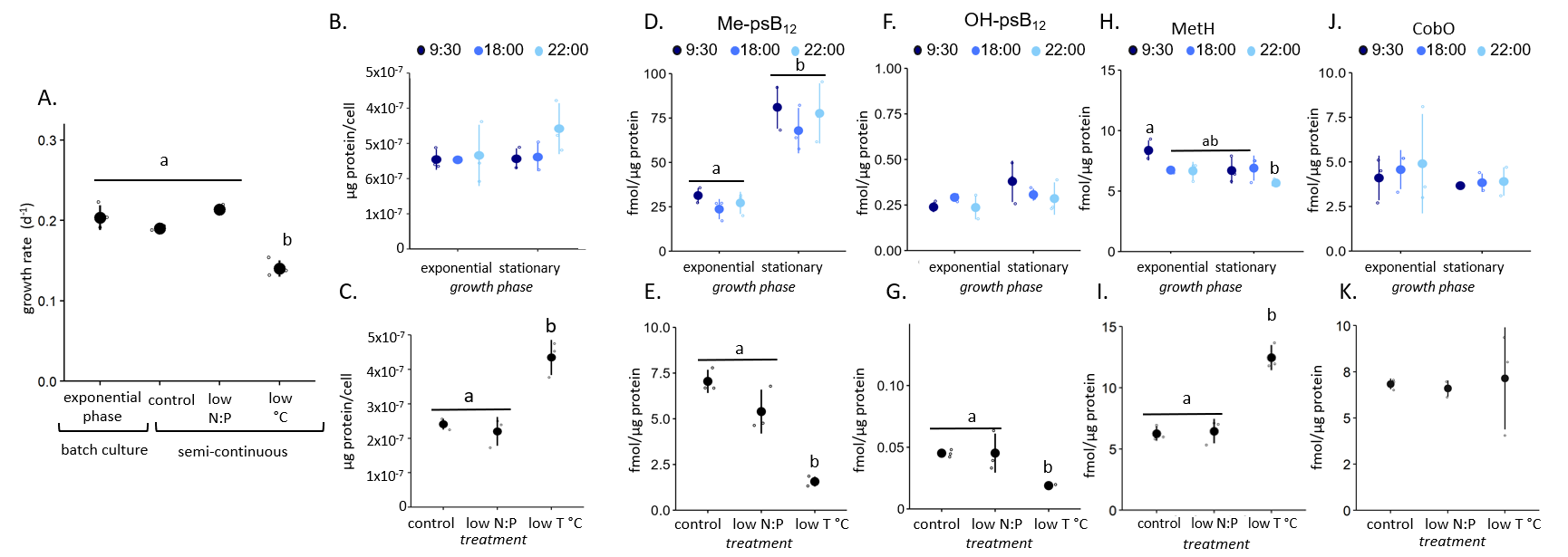

**Figure S3:** Changes in (A) growth rate (per day), (B,C) protein cellular content (fmol per µg protein), methyl- (D, E), hydroxy- (F, G) pseudocobalamin, and MetH (H, I) and CobO (J,K) (fmol per µgram total protein) in *Synechococcus* sp. WH 8102 under different experimental treatments grown in batch (B, D, F, H, J) and in semi-continuous cultures (C, E, G, I, K). Mean values are represented by solid circles, individual biological replicates (n=3) values by small open circles, and the standard deviation by lines. Different letters over data points indicate statistically significant differences (p-value < 0.05) between pairs of means based on the post-hoc Tukey's Test. Shared letters indicate no significant difference, if no letters are displayed then the differences between treatments were insignificant.

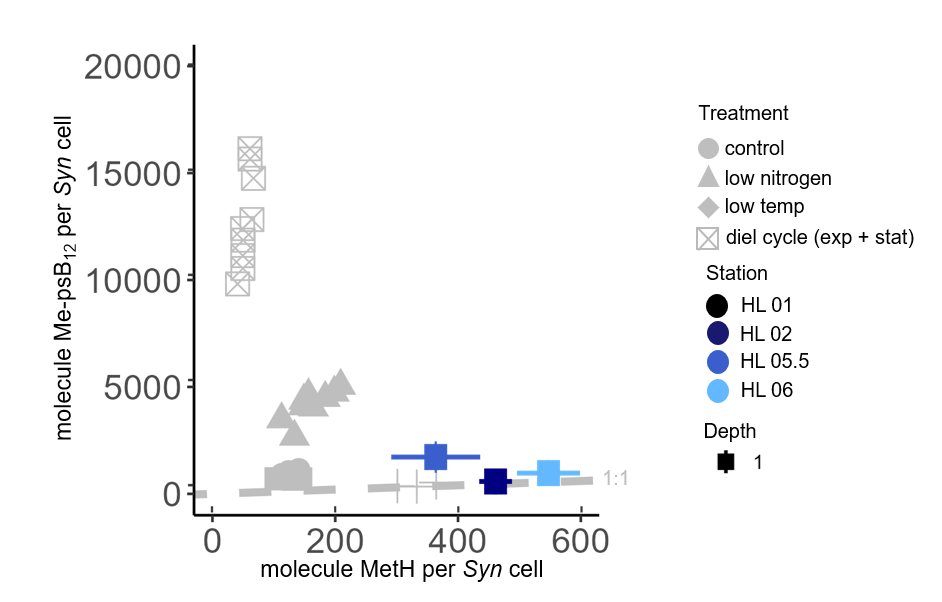

**Figure S4:** Molecule Me-psB_12_ per *Synechococcus* cell versus molecule of MetH per *Synechococcus* cell from culture samples across all treatments (grey points, biological replicates, n=3) and environmental data from stations on the Halifax Line, Northwest Atlantic Ocean, in Fall 2016 (blue points, lines represent technical replicates, n=1). Dotted line represents a 1:1 ratio of psB_12_ and MetH.

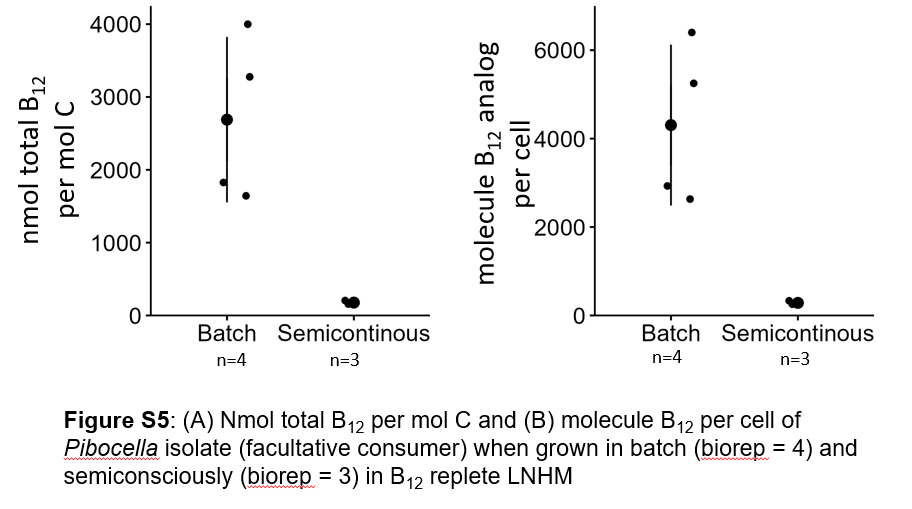

**Figure S5:** (A) Nano-mole total B_12_ per estimated mole carbon and (B) molecule B_12_ (Ado-, Me-, CN- and OH-B_12_ combined) per cell of *Pibocella* isolate when grown in batch (biological replicates = 4) and semiconsciously (biological replicates = 3) in B_12_ replete (737 pM) low nutrient heterotrophic media (LNHM).

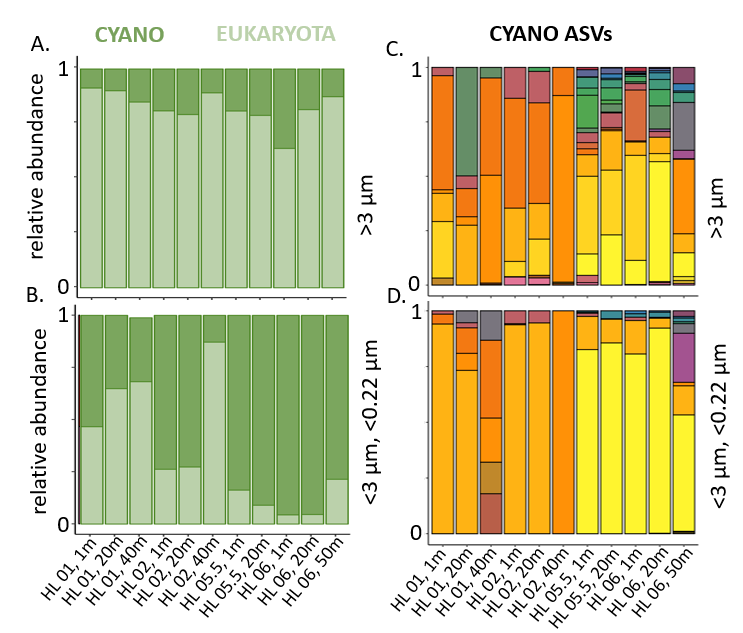

**Figure S6:** Relative abundance of 16S ASVs (chloroplast + cyanobacterial 16) for cyanobacteria (dark green) and eukaryotes (light green) from HL 01, HL 02, HL 05.5, and HL 06 (Fall 2016) on (A) large (>3 µm) and (B) small (<3 µm, >0.22 µm) size fractions. Relative abundance of all cyanobacterial ASVs (each unique color = 1 ASV) in the large (C) (>3 µm) and (D) small (<3 µm, >0.22 µm) size fractions. *Data from Robicheau et al, 2022.*

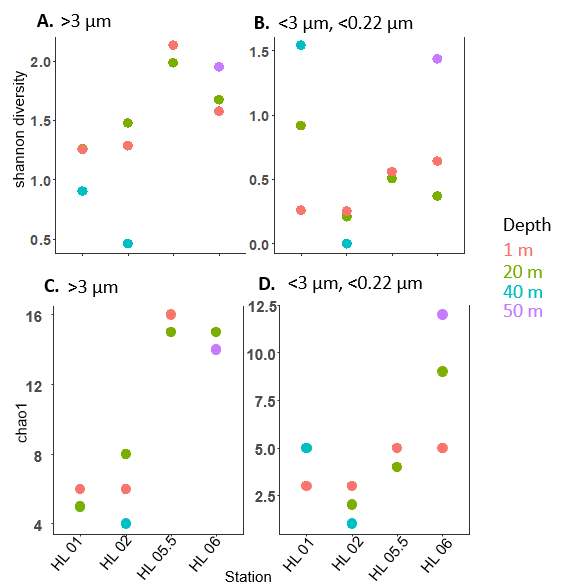

**Figure S7:** (A, B) Shannon diversity and (C,D) Chao1 (species richness) of 16S ASVs (chloroplast + cyanobacterial 16) for cyanobacteria and eukaryotes from HL 01, HL 02, HL 05.5, and HL 06 (Fall 2016) on (A, C) large (>3 µm) and (B, D) small (<3 µm, >0.22 µm) size fractions. *Data from Robicheau et al, 2022.*
